# Epithelia delimits glial apical polarity against mechanical shear to maintain glia-neuron - architecture

**DOI:** 10.1101/2022.12.26.521704

**Authors:** Cecilia G. Martin, James S. Bent, Aakanksha Singhvi

**Affiliations:** Division of Basic Sciences, Fred Hutchinson Cancer Research Center, Seattle, WA 98109; Current: Medical Scientist Training Program, University of Washington, Seattle, WA; Department of Biological Structure, University of Washington School of Medicine, WA 98195

**Keywords:** epithelia, glia, cell polarity, co-chaperone, sensory system, neuron cell shape

## Abstract

For an organ to maintain proper architecture and function, its different component cell-types must coordinate their cell-shapes with each other through life. While cell-intrinsic developmental mechanisms driving homotypic cell-cell coordination are known, how heterotypic cells collectively regulate cell-shape is less-clear. We report that, in a sense-organ, epithelial cells delimit and maintain polarity domains of contacting glia, and thereby, associated neuron shapes throughout life. Briefly, Hsp co-chaperone UNC-23/BAG2 keeps epithelial apical domains from deforming with animal movement. Epithelial apical domains stretch aberrantly and progressively in adult *unc-23* mutant animals, which in an FGFR-dependent manner, dislocates glial apical cytoskeleton proteins SMA-1/β_H_-Spectrin and actin. This alters glial apical polarity and cell shape, and concomitantly, associated neuron-ending shape. Notably, UNC-23 acts temporally at a developmental critical period to maintain glia-neuron shape in adults, and spatially within a defined anatomical zone. Lastly, intervention in either epithelia, glia or neuron ameliorate or phenocopy *unc-23* neural defects. Epi/endothelia resist mechanical stress and contact glia-neuron units across central/peripheral nervous systems and species, and all components of the identified molecular pathway are conserved and disease-relevant. Thus, we posit that analogous epithelia-glia mechanobiological coupling may broadly regulate glia-neuron shapes through animal life.

## INTRODUCTION

Our multicellular nervous systems are bustling with many chemical conversations. These allow component cells to coordinate their shape, organization and behavior with neighbors and function as a coherent collective. The sense-organ is an ideal platform to dissect such multi-cellular interactions. Each of our sense-organs, which mediates our ability to sense our environment and respond with meaningful behaviors, is a compartmentalized structural unit comprised of diverse cell types including sensory cells or neurons, specialized glia, and epithelial/endothelial cells (Ng et al., 2020). The shape of each sense-organ is exquisitely tuned to the modality sensed. However, while how each given cell type regulates its shape is studied in some detail, whether or how heterotypic cells in a sense-organ coordinate shape to build appropriate organ architecture and function through animal life is less explored.

Epithelia cells are an established model to study how homotypic cells communicate to regulate cell-shape. They coordinate both planar and apical-basal polarity axes with each other, driven by homotypic cell-cell signaling and maintained by the cell’s junctional complexes (Bosveld et al., 2018; Devenport, 2016). This derives the barrier functions of the skin epithelia as an organ, allowing it to resist mechanical shear and drive appropriate morphogenetic movements, tube formation, and create discrete polar cell-signaling microenvironments (Carvalho and Broday, 2020; Gibson and Perrimon, 2003; Horne-Badovinac and Bilder, 2005; Otani and Furuse, 2020; Pasti and Labouesse, 2014). Polarity of epi/endothelia also impacts neural development. In the peripheral nervous system (PNS), epithelia/like cells orient with specific cell-polarity in relation to neural tissue, and mechanical forces on developing neuroepithelia impact sensory placode development and neurogenesis (Phuong Le et al., 2022). However, if or how mechanical forces on epi/endothelia impact associated sense-organ shape or function in adults is not known.

Like epithelia, both major cell types of the nervous system, neurons and glia, are polarized. For neurons this polarity dictates directional transfer of information across the neural circuit. Neurons inherit molecular markers for apical-basal polarity from developing neuroepithelia and radial glia (Arimura and Kaibuchi, 2007; Barnes and Polleux, 2009; Bilder, 2001; Goldstein and Macara, 2007; Gotz and Huttner, 2005; Taverna et al., 2014). Components of these markers are repurposed to develop and maintain axon-dendrite process polarity (Cheng and Poo, 2012; Hapak et al., 2018; Lee et al., 2021; Szu-Yu Ho and Rasband, 2011; Wiggin et al., 2005).

A striking readout of this polarity is the neuron-receptive ending (NREs) on the dendrite process, a specialized sub-cellular structure through which the neuron receives inputs (Arimura and Kaibuchi, 2007; Higginbotham and Gleeson, 2007; Procko and Shaham, 2010; Shaham, 2010; Singhvi and Shaham, 2019). In the PNS, NREs sense environmental cues through specialized organelle shapes that track the specific modality sensed, such as modified photoreceptor cilia, or microvilli/stereocilia of tastebuds and inner ear (Carvalho and Broday, 2020; Elsaesser and Paysan, 2007; Gibson and Perrimon, 2003; Inglis et al., 2007; Manor and Kachar, 2008; Otani and Furuse, 2020; Pasti and Labouesse, 2014). Defects in PNS NRE shape are seen in patients with age-related sensorineural diseases or decline (e.g. deafness, blindness), or disorders of altered sensory acuity like Autism (Arimura and Kaibuchi, 2007; Elsaesser and Paysan, 2007; Estrada-Cuzcano et al., 2012; Higginbotham and Gleeson, 2007; Huang et al., 2020; Kremer et al., 2006; Lee *et al*., 2021; Riera and Dillin, 2016). Thus, development and maintenance of sensory NRE shape in a sense organ is key to maintaining sensory acuity with animal age. NREs contact either glia or glia-like epithelia in both CNS and PNS (Ray and Singhvi, 2021; Zuchero and Barres, 2015). In either settings, glia modulate NRE shape, and molecular mechanisms underlying this are only recently being uncovered (Allen and Eroglu, 2017; Christopherson et al., 2005; Chung et al., 2015; Haber et al., 2006; Murai et al., 2003; Procko and Shaham, 2010; Singhvi and Shaham, 2019). Glia inherit polarity from neuroepithelial stem cell progenitors and exhibit elaborate and diverse cell shapes, presumably tracking function (Cobo et al., 2020). Besides NREs, glia also contact epi/endothelia, including at sense-organ epithelia, skin epithelia, blood vessel endothelia, or meninges (Abbott, 2002; Cobo *et al*., 2020; Derk et al., 2021; Gulbransen and Sharkey, 2012; Lakkaraju et al., 2020; Langen et al., 2019). In the limited instances examined, post-development glia exhibit apical-basal polarity signatures, with apical regions apposing neurons and basolateral domains proximal to epi/endothelia (Derouiche et al., 2012; Low et al., 2019; May-Simera and Kelley, 2012; Salzer, 2003; Silies and Klambt, 2011). How glia coordinate and maintain their polarized cell-shape and orientation with both NREs and endothelia/epithelia is poorly understood. Further, while glia dictate epithelial barrier functions, it is unclear if epithelia reciprocally dictate glial properties.

To study how coordinate glia-neuron cellular architectures are maintained, we turned to *C. elegans*, a powerful and genetically tractable experimental model to investigate glia cell-biology and glia-neuron interactions at molecular mechanistic and single-cell resolution (Ray and Singhvi, 2021; Singhvi et al., 2016; Singhvi and Shaham, 2019; Wallace et al., 2016). The animal’s nervous system comprises 50 ectodermal-lineage derived glia (like PNS glia or astrocytes) and 300 neurons. Prior work by us and others has shown that *C. elegans* glia regulate associated NRE shape and functions, with impact on animal sensory behavior and memory (Oikonomou and Shaham, 2011; Raiders et al., 2021b; Ray and Singhvi, 2021; Shaham, 2010; Singhvi and Shaham, 2019).

One *C. elegans* glia is the amphid sheath glia (AMsh), part of the animal’s major sense organ called the amphid. It interacts with twelve sensory amphid NREs at its anterior apical domain, and with epithelial cells at its basolateral domain (Low *et al*., 2019; Perkins et al., 1986; Ward et al., 1975) (SR, personal communication). One contacting NRE is that of the animal’s major thermosensory neuron, AFD. The AFD-NRE comprises ∼45 microvilli of ∼5-12µm length (henceforth, AFD-NRE-m) and a single cilium ∼1-2µm length (noted as AFD-NRE-c) (Figure 1A) (Doroquez et al., 2014; Perkins *et al*., 1986; Singhvi *et al*., 2016). We and others have previously identified both AMsh glial and cell-intrinsic regulators of AFD-NRE-m shape, many with age-dependent defects (Huang *et al*., 2020; Inada et al., 2006; Lewis and Hodgkin, 1977; Perkins *et al*., 1986; Raiders et al., 2021a; Ray and Singhvi, 2021; Satterlee et al., 2001; Singhvi *et al*., 2016; Singhvi and Shaham, 2019; Wallace *et al*., 2016). All glial regulators are evolutionarily conserved, and their loss lead consequently to animal thermosensory behavior deficits (Huang *et al*., 2020; Raiders *et al*., 2021a; Singhvi *et al*., 2016; Wallace *et al*., 2016). Thus, AMsh glia-AFD neuron pair is a facile setting to interrogate how glia-neuron units establish and maintain cell-shape through animal life, at exquisite single-cell resolution.

**Figure 1.**
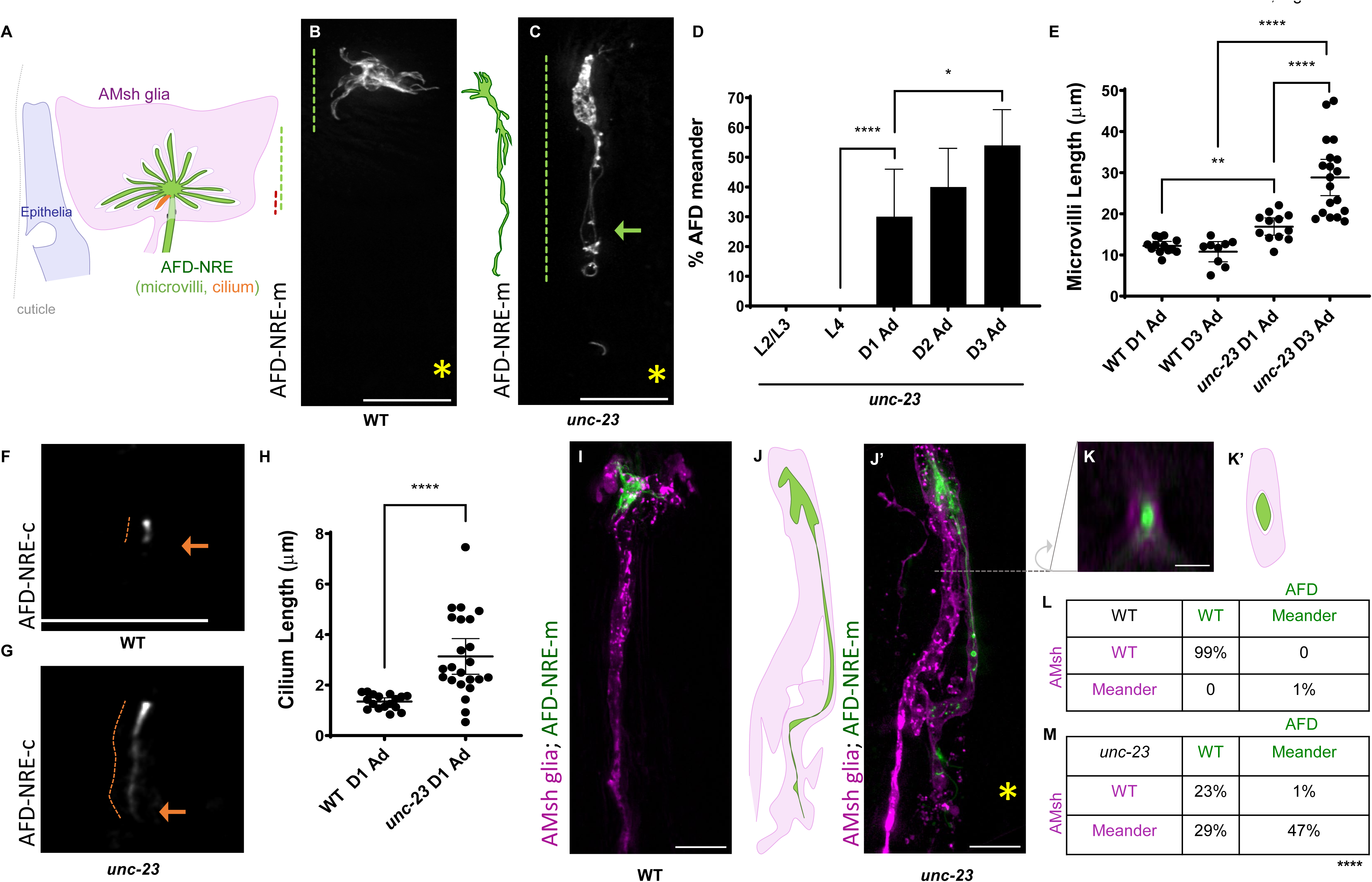
UNC-23 regulates amphid sense-organ architecture. (**A**) Epithelia (blue) appose AMsh glia:AFD neuron unit. AMsh glia (magenta) ensheathes the composite AFD-NRE (green microvilli, single red cilium). (B-M) All data refers to *unc-23(e25)* (**B, C**) Fluorescence images of AFD-NRE microvilli labeled with *nsIs645* in (B) WT Day 1 adults and (C) *unc-23* Day 1 adults. Region occupied is denoted by dotted green line. Arrow = region of overgrowth, asterix = site of first pharyngeal bulb. Scale bar, 8μm. (**D**) AFD-NRE-m meandering is age-dependent. L2/L3 (N=43), L4 (N=55), D1 Ad (N=30), D2 Ad (N=62), and D3 Ad (N=72) where L= larval stage, Ad= adult, and D= day. Data plotted as population sum ± 95% CI. Two-sided Fisher’s Exact Test * p=0.0308 and **** p<0.0001. (**E**) Average AFD-NRE microvilli length (μm) in WT Day 1 adults (mean= 12.23 ± 1.08 95% CI, N=13), WT Day 3 adults (mean= 10.81 ± 2.467 95% CI, N=9), *unc-23* Day 1 adults (mean= 16.91 ± 2.11 95% CI, N=12), and *unc-23* Day 3 adults (mean=28.86 ± 4.41 95% CI, N=19). Data plotted as mean ± 95% CI. Dunnett’s T3 Multiple Comparisons Test ** p=0.0017, **** p<0.0001. (**F, G**) Fluorescence images of AFD-NRE cilium in WT (F) and *unc-23* (G) Day 1 adults. Arrow points to cilium. Overgrowth in mutant denoted by dotted red line. Scale bar, 8μm. (**H**) Random sampling AFD-NRE cilium length (μm) for WT (mean= 1.349 ± 0.15 95% CI, N=17) and *unc-23* (mean= 3.138 ± 0.705 95% CI, N=23) Day 1 adults. Data plotted as mean ± 95% CI. Welch’s t-test **** p<0.0001. (**I**) AMsh (magenta) and AFD-NRE (green) process in WT. Scale bar, 8μm. (**J, J’**) Schematic and image of AMsh:AFD in *unc-23* mutants. Asterix = site of anterior bulb. Scale bar, 8μm. (**K-K’**) Orthogonal XZ section as schematic and image of AMsh:AFD-NRE at dotted line in J’. Asterix = site of anterior bulb. Scale bar, 2μm. (**L, M**) Correlation analysis of AFD-NRE microvilli meander phenotype with AMsh glial meander phenotype in WT (L) and *unc-23* mutant animals (M). Two-sided Fisher’s Exact Test **** p<0.0001. N=60 for wild type and N=73 for *unc-23*.

We report here that mechanobiological coupling of epithelia and glia against external shear impacts neuron shape through animal life. By focused inquiry of the AMsh glia-AFD neuron pair, we also uncover the underlying molecular mechanism for this tricellular interaction. Briefly, the Hsp co-chaperone UNC-23/BAG2 acts in epithelial cells (the animal’s skin/ epidermis) to delimit their apical domain boundary under mechanical shear stress from animal locomotion, and its loss leads to aberrant expansion of epithelial apical domains. In an EGL-15/FGFR dependent manner, this leads to loss of the apical actin-associated cytoskeletal component SMA-1/β-Spectrin in apposed AMsh glia and consequent disorganization of glial F-actin. Expectedly, both *unc-23* and *sma-1* mutant animals exhibits defects in glial cell-shape and apical-domain polarity. In turn, this causes overgrowth of glia-associated NREs that contact glial apical membranes, including AFD-NRE. Surprisingly, while sensorineural defects present progressively only in adult animals, we find that UNC-23 functions in a critical larval developmental time window, to subsequently maintain glia-neuron shape and sense-organ architecture in adults. Furthermore, we find that UNC-23 regulates glia only within an anatomical region and impacts different glia differently. This, this previously unappreciated epithelia-glia mechanobiological coupling occurs with striking cellular heterogeneity . Finally, interventions in either epithelia (block mechanical shear), glia (block secretome) or neurons (increase activity) can all suppress *unc-23* neural defects. Thus, this study reveals a previously unappreciated role for heterotypic epithelial cells in maintaining architecture of sense-organ glia-neuron units through life. Conservation of all components identified suggests that analogous tricellular communication may broadly govern glia-neuron shape maintenance and aging.

## RESULTS

### A mutation with progressive defects in AFD-NRE and associated AMsh glia shape

To define regulators of NRE shape, we performed a forward genetic screen (10,800 F2 animals /2.5x genome) for adult mutant animals with defective AFD-NRE-m. One mutant lesion recovered, *ns344*, exhibited dramatic overgrowth of AFD-NRE-m with 34% penetrance, a phenotype we term “meander” (Figure 1B-D). These animals have overgrowth of multiple microvilli, and the phenotype is temperature-sensitive (Figure S1A). Neurite length measurements revealed a 1.4x elongation in AFD-NRE-m length (Figure 1D) (mean length: *ns344*= 16.9µm ± 3.3µm compared to *wild type* = 12.2µm ± 1.8µm). To control for pleiotropic defects in animal size in the mutant, we normalized analyses to either pharynx or buccal cavity lengths, and again noted a 1.5x increase in AFD-NRE-m length (Figure S1B) (mean length: *ns344*= 18.1µm ± 3.6µm compared to *wild type* = 12.2µm ± 1.8µm). A smaller yet persistent fraction of *ns344* animals also have missing microvilli, a phenotype we term “collapsed” (Figure S1C-S1E). Genetic analyses revealed “meander” as the true recessive phenotype (see Methods), which we focused on for subsequent analyses. To examine AFD-NRE cilia, we examined transgenic animals that had AFD-NRE-c marked by cell-specific tagging of the ciliary protein DYF-11/TRAF31B1 (Figure 1F-H)(Bacaj et al., 2008a; Kunitomo and Iino, 2008; Li et al., 2008; Raiders *et al*., 2021a; Starich et al., 1995). While mean AFD-NRE-c length was 1.35µm ± 0.29µm in wild type animals, matching prior reports (Doroquez *et al*., 2014; Perkins *et al*., 1986), those in *ns344* mutant animals were 3.12µm ± 1.63µm (2.3x increase) (Figure 1H). Thus, the *ns344* lesion perturbs both AFD-NRE microvilli and cilia.

Since AFD-NRE-m shape shows age-dependent changes (Huang *et al*., 2020), we examined *ns344* AFD-NRE defects longitudinally. AFD-NRE and glia shape of *ns344* mutant animals were unaltered in larva and defects presented primarily in Day 1 adult animals (Figure 1D, 1E). This was progressive, and in Day 3 adults, we observed a 2.7x increase in normalized microvilli lengths (Figure 1D, 1E) (mean length: *ns344*= 28.9µm ± 9.1µm compared to *wild type* = 10.8µm ± 3.2µm). Analyses were capped at Day 3 since mutant animals also had reduced lifespans (5.02 ± 0.29 day mean lifespan, compared to *wild type* 14.91 ± 0.46 day mean lifespan) (Figure S1F). Thus in *ns344* mutant animals, both AMsh glia and AFD-NRE do not maintain proper cell-shape in adult animals, and reduced organismal lifespan.

Since AFD-NRE are embedded in AMsh glia (Ward *et al*., 1975), we wondered if meandering AFD-NRE breach AMsh glial ensheathment. Examination of AMsh glia in mutant animals revealed that the glial anterior endings also exhibit “meander” shape defects (Figure 1I-K’). Correlative analysis of AMsh glia and AFD-NRE-m co-labeled with different fluorescent reporters showed that elongated AFD-NRE-m remain ensheathed by meandering AMsh processes (Figure 1L, 1M). Thus, *ns344* causes concomitant defects in both glia and NRE shape.

### *ns344* is a mutation in the Hsp co-chaperone, UNC-23/BAG2

We mapped *ns344* as a lesion in *unc-23* (UNC-23B^E238K^), identical to the reference *unc-23(e25)* allele isolated nearly 50 years ago in the first genetic dissection screen for *C. elegans* behavior and muscle tone (Brenner, 1974; Meissner et al., 2009; Rahmani et al., 2015; Waterston et al., 1980). Both *unc-23(e25)* and *unc-23(e324)*, a truncation allele, phenocopied *unc-23(ns344)* AFD-NRE-m shape defects (Figure 2A, 2B, S2A) and have detachment of anterior head musculature. Surprisingly, the deletion allele *unc-23(ok1408)* (Papsdorf et al., 2014) exhibited a significantly weaker AFD-NRE-m phenotype (Figure 2B). *in silico* breakpoint analyses of *ok1408* locus suggested that exon-skipping may generate a truncated in-frame transcript with the BAG domain, which was confirmed by RT-PCR analyses (Figure 2A, S2B). Thus, *ok1408* allele is a molecular hypomorph, which explains its weaker phenotype.

**Figure 2.**
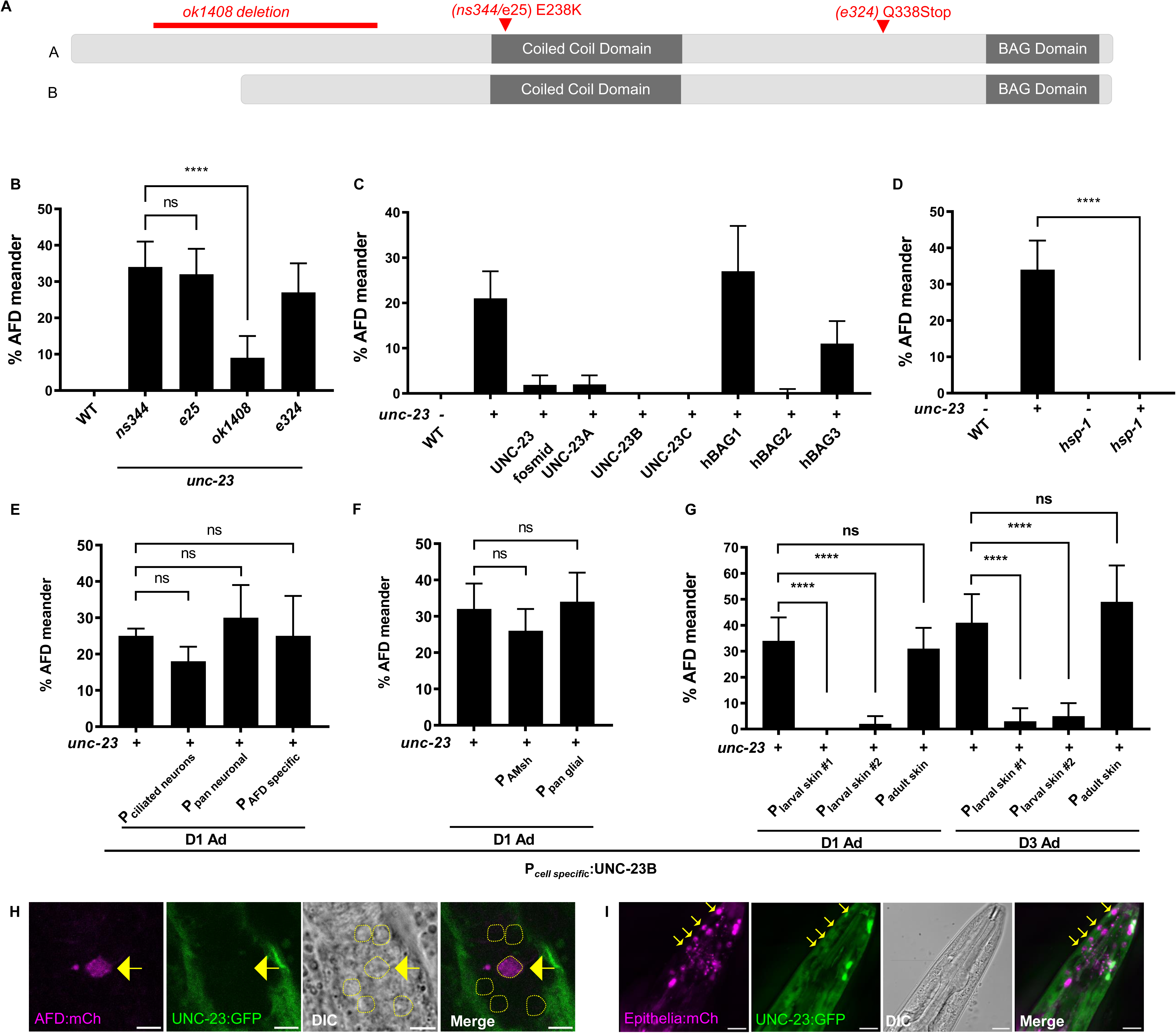
UNC-23/BAG2 acts in epithelia to maintain neuron shape in adults. (**A**) Schematic of protein domains and lesions in UNC-23^A^ and UNC-23^B^ isoforms, as noted. (**B-G**) AFD-NRE marked by *nsIs228* (B-F) or *nsIs645* (G) with % AFD-NRE meander in genotypes noted. Data is plotted as population sum ± 95% CI and a Two-Sided Fisher’s Exact Test was used for all statistics. (B) N≥108 for each genotypes. ns p=0.9045, **** p<0.0001. (C) N≥67 for all genotypes. **** p<0.0001, ** p=0.0050. (D) N≥90 for all genotypes. **** p<0.0001. (**E, F, G**) ≥2 transgenic extra-chromosomal lines were used (E) P*_dyf-11_*, P*_rab-3_*, and P*_srtx-1b_* drove ciliated neuron, pan-neuronal, and AFD specific UNC-23B expression respectively. N≥60 for all genotypes scored as Day 1 adults. ns p=0.3309 (P_dyf-11_), ns p=0.0865 (P*_rab-3_*), and ns p=0.6026 (P*_srtx-1b_*). (F) P*_F53F4.13_* and P*_mir-228_* drove AMsh specific and pan-glial UNC-23B expression respectively. N≥134 for all genotypes scored as Day 1 adults. ns p=0.2127 (P_F53F4.13_) and ns p=0.9022 (P_mir-228_). (G) P*_dpy-7_* and P*_rol-6_* drove expression of UNC-23B in larval epithelia and P*_col-19_* drove UNC-23B in adult epithelia. N≥122 and N≥45 for all genotypes scored as Day 1 adults and Day 3 adults respectively. **** p<0.0001, ns p=0.5875 (P*_col-19_* D1 Ad), and ns p=0.4591 (P_col-19_ D3 Ad). (**H)** Representative fluorescence images and DIC of AFD neuron cell body (magenta, arrow) and UNC-23:GFP (green). Dotted outlines mark other neuron cell bodies on DIC, Merge panels show absence of UNC-23:GFP (green) in AFD neuron (magenta). Scale bar, 5μm. (**I**) Representative two-color fluorescence and DIC image of UNC-23:GFP (green) and epithelial nuclei (magenta). Merge panel shows UNC-23 expresses in epithelia. Arrows denote nuclei position in GFP+ cells. Scale Bar, 15μm.

UNC-23 is a member of the BAG family proteins, nucleotide exchange factors that facilitate substrate cycling and chaperone-assisted protein folding through the Hsp70/Hsc70 machinery (Arndt et al., 2007; Hoppe and Cohen, 2020; Kabbage and Dickman, 2008; Mayer and Bukau, 2005; Takayama et al., 1999). Humans have 6 BAG protein sub-families (BAG1-6). UNC-23 aligns closest to BAG2 and is also proposed to represent composite characteristics of BAG2, 3, and 4 (Kabbage and Dickman, 2008; Papsdorf *et al*., 2014; Takayama *et al*., 1999). The *unc-23* locus has three predicted isoforms (UNC-23^A/B/C^) that differ in start codon site (Figure S2A). AFD-NRE defects of mutant animals were completely rescued by expression of either *unc-23* fosmid or cDNA encoding any of the three predicted isoforms under a pan-cellular promoter (Figure 2C), indicating that shared isoform function drives regulation of AFD-NRE shape. We also engineered “humanized” transgenic animals that express either human BAG1, hBAG2, or hBAG3 globally. Expression of hBAG2 completely rescued AFD-NRE-m defects of *unc-23(e25)* mutant animals, the closely related hBAG3 rescued partially, and hBAG1 does not rescue (Figure 2C). Thus, *ns344* lesion in UNC-23/BAG2 regulates AFD-NRE shape.

### UNC-23 acts with HSP-1/Hsp70 to regulate AFD-NRE

BAG2 family members contain a C-terminal BAG/BNB domain that binds Hsp chaperones, and an N-terminal region with a coiled-coil motif with proposed regulatory function (Figure 2A) (Papsdorf *et al*., 2014; Papsdorf et al., 2019). The *e25/ns344* lesion (UNC-23^E238K^) maps to the N-terminal region, suggesting two plausible molecular models of action. Either the E238K Lys creates an aberrant ubiquitination site leading to UNC-23 degradation, or this charge-altering mutation caused protein misfolding/function which impairs binding of either a regulator or Hsp. Three lines of evidence favor the latter model. One, *in silico* fold-prediction of UNC-23B, UNC-23B^E238K^, UNC-23B^E238R^, and UNC23B^E238D^ suggest that a positively charged residue at E238 will unfold the BAG domain (Fig S2C), consistent with previous CD spectroscopy studies of thermal stability (Papsdorf *et al*., 2014; Papsdorf *et al*., 2019). Two, UNC-23^E238R^, which bears a similarly charged mutation that is not ubiquitin-prone is unable to rescue the mutant phenotype (Figure S2D). Three, analysis of *hsp-1*. A mutation in the *C. elegans* HSP-1/Hsp70 ortholog, *hsp-1(ra807)*, was identified as a suppressor of *unc-23* musculature defects and has compensatory alteration of protein interaction site (Papsdorf *et al*., 2014; Papsdorf *et al*., 2019; Rahmani *et al*., 2015). We find that it also suppressed *unc-23* AFD-NRE defect (Figure 2D). Interestingly, *hsp-1(ra807)* exhibit AFD-NRE collapse defects like *unc-23* lesions, and this was not further enhanced in the double mutant (Figure S2E). Alluding to the specificity of UNC-23: HSP-1 interaction, loss of two other Hsp70 family genes (*hsp-3, hsp-70)* did not exhibit equivalent degree of AFD-NRE defects or suppression in wild type or *unc-23* mutant animals, respectively (Figure S2F, S2G). Thus, we infer that AFD-NRE shape defects are due to altered UNC-23:HSP-1 binding and activity.

### UNC-23 acts developmentally in skin epithelia to regulate AFD-NRE shape in adults

UNC-23 expresses in many tissues (Papsdorf *et al*., 2014; Rahmani *et al*., 2015). We first asked if UNC-23 acted cell-autonomously in AFD to regulate its NRE shape. Surprisingly, there was no rescue of *unc-23(e25)* defects when UNC-23B was expressed under either of three neuronal promoters (ciliated-neuron-specific *P_dyf-11_*; pan-neuronal *P_rab-3_*, and AFD-specific *P_srtx-1_*) (Figure 2E). Consistent with this, an *unc-23:gfp* translational reporter does not express in AFD (Figure 2H). Since AMsh glia ensheath AFD-NRE and modulate its shape (Raiders *et al*., 2021a; Singhvi *et al*., 2016; Wallace *et al*., 2016), we next asked if UNC-23 functions in glia. This was also not the case: UNC-23B expressed under either AMsh specific (P*_F53F4.13_*) or pan-glial promoter (P*_mir-228_*) failed to rescue *unc-23* mutant defects (Figure 2F). Thus, UNC-23B acts from a non-neural cell to regulate AFD-NRE shape.

Since *unc-23* lesions also exhibit progressive detachment of anterior head musculature, we next asked if neural defects were secondary to muscle defects. However, not all mutations that cause muscle detachment co-present with AFD-NRE defects (*mua-10, vab-10/Spectraplakin*) (Plenefisch et al., 2000) (Figure S2H). In corollary, not all mutations or genetic backgrounds that cause AFD-NRE overgrowth, or act with *unc-23*, impact muscle attachment (see below). We infer that NRE and muscle defects are not causally linked in *unc-23* mutants.

Finally, we noted that at the anterior tip, AMsh glia contact both NREs and the animal’s skin epithelia cells (also called epidermis or hypodermis), and epithelia express UNC-23 (Figure 2I) (Altun et al., 2002; Brenner, 1974; Rahmani *et al*., 2015; Waterston *et al*., 1980). Remarkably, UNC-23B expression in skin epithelia under either of two promoters (P*_dpy-7_* and P*_rol-6_*) completely rescued mutant AFD-NRE-m defects (Figure 2G). Thus, UNC-23/BAG2 in the overlying epithelia maintains AMsh glia-AFD neuron unit shape. Of note, while both promoters rescue neural defects similarly and completely, they rescue muscle defects differently, bolstering the notion that muscle and neural defects are not causally linked in *unc-23* mutants (Figure S2I).

Curiously, both P*_dpy-7_* and P_rol-6_ epithelial promoters are active in larval development and switch off in adults, with P*_rol-6_* expressing specifically during larval molts in the major epidermis and excluded from the specialized lateral hypodermis/seam cells (Kramer and Johnson, 1993; Park and Kramer, 1994; Sassi et al., 2005). This temporality was surprising since we find that *unc-23* mutant neural defects manifest primarily in adults and are progressive with age, with larval animals being normal (Figure 1D). Further, expression of UNC-23B under either promoter not only rescues Day 1 adult defects, but also persists into Day 3 adults (Figure 2G). To test if UNC-23 acts in adults, and the persistent rescue reflects larval-expressed UNC-23B protein perduring into adults, we also expressed UNC-23B under the adult -specific promoter P*_col-19_*. This could not rescue *unc-23(e25)* mutant AFD-NRE defects (Figure 2G). Thus, UNC-23 acts in epithelia of molting larva (likely L4-adult molt), and its loss in this spatiotemporal developmental window results in progressive neural defects in post-molt adults.

### Epithelial UNC-23 delimits apical domains in both epithelia and glia

How might epithelial UNC-23 regulate neural defects? Prior work has reported that liquid culture growth suppresses muscle defects of *unc-23(e25)* (Rahmani *et al*., 2015). Since our data decoupled muscle and neural defects, we asked if liquid growth independently suppressed neural defects, and found that it does (Figure 3A). Further, mutations in the myosin heavy chain gene *unc-54,* which render the animal immobile, also suppressed *unc-23(e25)* AFD-NRE-m defects (Figure 3A). Thus, UNC-23 allows the animal to withstand mechanical stress while crawling on solid media, and loss of this function impacts AFD-NRE shape.

**Figure 3.**
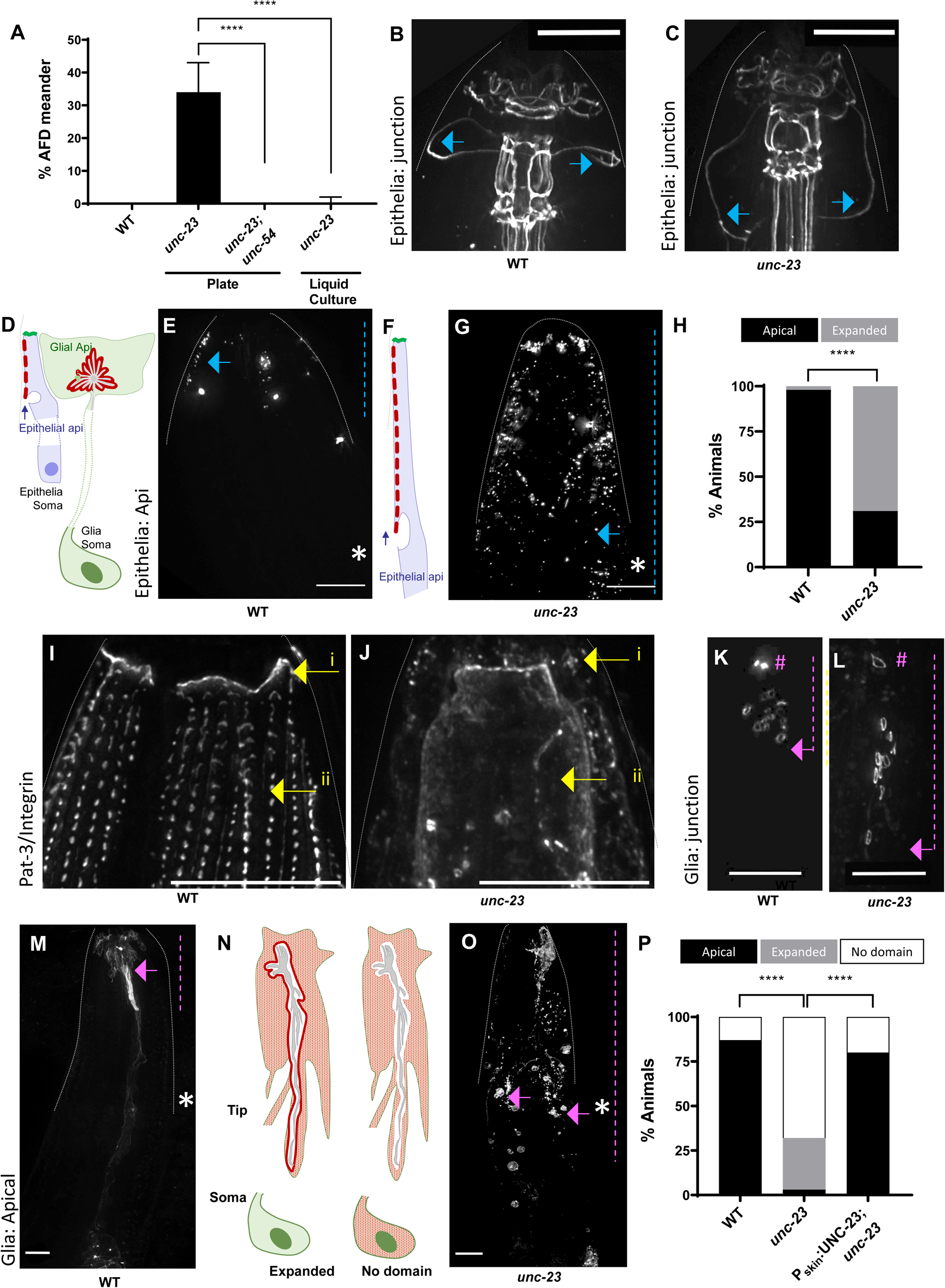
UNC-23 delimits epithelial and AMsh glial apical domains from mechanical stress. All data based on *unc-23(e25)* allele and anterior nose-tip on top in all images (dotted white live) (**A**) *nsIs645* AFD-NRE microvilli marker. % AFD-NRE meander in liquid culture [2 biological replicates, WT (=112), *unc-23* (N=186)] or *unc-54(e190)* immobilized animals (N≥112). Data plotted as population sum ± 95% CI. Two-sided Fisher’s Exact Test **** p<0.0001. (**B, C**) Epithelial tight junction marker DLG-1:GFP. Blue arrow = hyp4:hyp5 epithelial cell junctions in wild type, displaced posteriorly in *unc-23* mutant animals. Scale bar, 8μm. (**D-G**) Schematic and representative fluorescence images of epithelial apical domain marker in wildtype (D, E) and *unc-23* mutant animals (F, G). Apical domain marker (blue arrow), with domain boundary (dotted blue line) respective to midpoint of anterior pharyngeal bulb (asterix). Scale bar, 8μm. (**H**). Quantification of apical restriction: wild type (N=62), *unc-23* (N=56). Two-sided Fisher’s Exact Test **** p<0.0001. (**I, J**) Fluorescence images of anterior head region PAT-3:mNeonGreen in WT (I) and *unc-23(e25)* (J). Loss of PAT:GFP staining (yellow arrows i, ii). Scale bar, 8μm. (**K, L**). Fluorescence images of AMsh tight junctions in wild type (K) and *unc-23* (L), with junctional zone denoted (pink line). Pink arrow = posteriorly displaced glia:neuron junctions in *unc-23* mutants. # = normal AMsh:AMso junction in wild type and *unc-23* mutants. Scale bar, 8μm. (**M-O**) Schematic and representative fluorescence images of glial apical domain marker in wildtype (M) and *unc-23* mutant animals (N, O). Apical domain (pink arrow) with boundary (pink dotted line) respective to mid-point of anterior pharyngeal bulb (asterix). Scale bar, 8μm. (**P**) Quantification of glial apical domain boundary enrichment in wild type (N=159), *unc-23* (N=92), or *unc-23;* P*_epithelia_*:UNC-23B (N=49). Two-sided Fisher’s Exact Test **** p<0.0001.

Epithelia withstand mechanical tension by coordinating apical-basal polarity and junctions across each other to function as a sheet. We therefore evaluated these cellular features in epithelia of *unc-23* mutant animals. First, we examined cell:cell junction DAC complex component DLG-1/*Discs Large* i*n vivo* (McMahon et al., 2001). DLG-1 maintained localization to junctional boundaries in *unc-23* mutants, suggesting that its loss does not underlie mutant neural defects (Figure 3B, C). However, we observed that junctions between specific epithelia cells (hyp3-hyp6) in the anterior head region were mis-localized posteriorly (Figure 3B-C, arrow). This led us to wonder if apical-basal domain boundaries were affected. To examine this, we engineered genetic reagents to mark epithelial apical and basolateral domains by adapting previously described apical-basal markers for glia (Low *et al*., 2019). We noted significant expansion of apical domain in *unc-23* mutant epithelia (Figure 3D-H) and posterior displacement of basolateral domain markers (Figure S3A-B”, green staining). This could reflect either inability to specify basolateral domain identity, maintain apical-basal sorting, or cell stretching under mechanical shear. The thin epithelia in the head have minimal cytoplasmic milieu, which is exaggerated in *unc-23(e25)* (Rahmani *et al*., 2015; Waterston *et al*., 1980). This precludes analysis of apical-basal marker overlay in epithelia by fluorescence light microscopy reporters. So, we probed integrin receptors, which decorate basolateral domains to tether it to ECM (Lee and Streuli, 2014). Loss-of function mutation in INA-1/α-Integrin or RNAi knock-down of PAT-3/β-integrin, or the ECM component UNC-52/Perlecan, neither phenocopied nor suppressed *unc-23* mutant animals (Figure S3C, S3D). Consistent with our basolateral domain marker phenotype and similar to UNC-52/Perlecan (Rahmani *et al*., 2015), we found that INA-1 and PAT-3 staining was lost and/or disorganized in the anterior head region of the animal (Figure S3E-F, 3I-J). We infer that stretching of the apical domain under mechanical shear stress posteriorly displaces epithelial basolateral domains in relation to animal body plan.

We similarly examined glial apical-basal polarity in *unc-23* animals. AMsh glia apical domains contact embedded NREs like those of AFD, the DAC marker AJM-1 localizes to AMsh: NRE and AMsh glia: AMso glia contacts, and glial basolateral domain apposes epithelia (Altun *et al*., 2002; Low *et al*., 2019; Pasti and Labouesse, 2014) (CM, SR and AS, personal communication). In *unc-23* mutant animals bearing an AMsh glia-specific fluorescently tagged AJM-1, AMsh: AMso cell junctions were unaffected, and AMsh: NRE remained intact albeit posteriorly displaced, similar to junctional markers in epithelia (Figure 3K, 3L).

Expression of AMsh glia-specific apical markers, on the other hand, revealed striking defects. While 87% *wild type* animals showed apical restriction of a transgenic apical marker, only 9% of *unc-23* mutant animals did so, with 52% animals having no discernable apical domain restriction, and an additional 39% animals with expanded apical domains (Figure 3M-3P). Further, this aberrant glial apical expansion, like AFD-NRE defects, was rescued by expression of UNC-23B under P*_dpy-7_* epithelia-specific promoter (Figure 3P). Given the similar phenotypes of epithelia and glia, we also double-labeled epithelial basolateral domains with glial apical domains and found that they do not overlap in either wild type or *unc-23* mutant animals (Figure S3A-B’’). Taken together, we infer that glial apical polarity tracks epithelial apical polarity i.e. UNC-23’s regulation of epithelial apical domains causes defects in glial apical domain limits and cell-polarity, and thereby NRE shape.

### UNC-23 acts with glial β_H_-spectrin to regulate AFD-NRE shape

How does UNC-23 induce glial cell-polarity and shape defects? Spectrins and spectraplakins are large cytoskeletal proteins that maintain cell shape by crosslinking F-actin networks to itself, the cell membrane or microtubule networks (Cusseddu et al., 2021; Deng et al., 2020). Spectrins are obligate dimers of α-and β-subunits. The *C. elegans* genome encodes one α-spectrin (SPC-1), two β-spectrins (apical-domain specific SMA-1 and basolateral-domain enriched UNC-70), and one Spectraplakin (VAB-10), and UNC-70/β_G_-spectrin protects axon process fidelity (Jia et al., 2019; Moorthy et al., 2000). Given *unc-23’s* cell-biology defects led us to explore if mutations in any of these genes also phenocopied its AFD-NRE dendrite defects.

Loss of neither VAB-10/Spectraplankin nor UNC-70/β_G_-spectrin phenocopy *unc-23* AFD-NRE shape defects, suggesting distinct cell-biology (Figure S2H, 4A). SPC-1/ α-spectrin expresses in many cells including epithelia, neurons, and glia (Jia *et al*., 2019; Norman and Moerman, 2002), but analysis was precluded by animal lethality. While a single-copy GFP:SPC-1 CRISPR-transgene (Jia *et al*., 2019) showed extensive disorganization in *unc-23* mutants, its ubiquitous expression precluded conclusive inference (Figure S4A, A’). Strikingly, however, three mutant alleles in the apical SMA-1/β_V_/β_H_-spectrin phenocopied *unc-23(e25)* AFD-NRE defects (allele *e30 =* 20% meanders, 27% AFD-NRE collapse, n= 88) (Figure 4A, S4B-D). Measurement of AFD-NRE-m lengths in *sma-1(e30)* mutant animals revealed a 1.6x overgrowth (mean length: *sma-1*(*e30)=*19.2µm ± 4.5µm; compared to *wild type* = 12.2µm ± 1.8µm and *unc-23(e25)* = 16.9µm ± 3.3µm) (Figure 4B). When normalized for animal body size, this reflects a 2.4x overgrowth (mean length: *sma-1*(*e30)=* 29.3µm ± 6.9µm; compared to *wild type* = 12.2µm ± 1.8µm and *unc-23(e25)* = 18.1µm ± 3.6µm) (Figure S4E). Like in *unc-23(e25)* animals, AMsh glial apical-basal polarity was also disrupted in *sma-1* mutant animals (Figure 4C-E) and their neural defects trend progressively, although not statistically significant (Figure 4B, S4E). Unlike *unc-23(e25)* animals, *sma-1* mutants do not display muscle detachment (data not shown), again decoupling AFD-NRE growth from muscle detachment defects.

**Figure 4.**
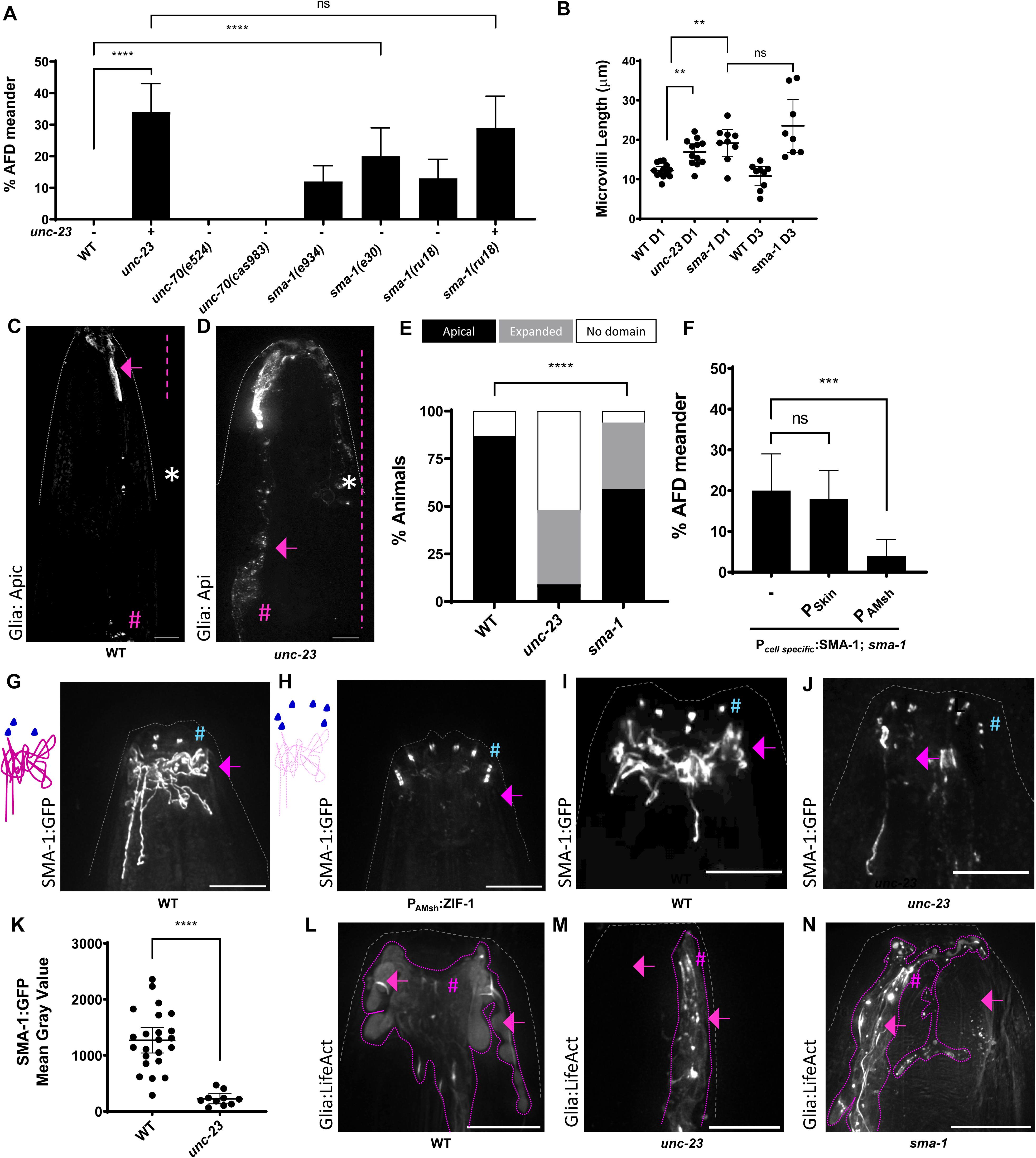
UNC-23 regulates AFD-NRE through glial β_H_-Spectrin. Analyses on *unc-23(e25)* and *sma-1(e30)* alleles throughout unless otherwise noted, *nsIs645* AFD-NRE marker. Anterior nose-tip to top with animal outline (dotted gray). (**A**) % AFD-NRE meander. N≥88 except *unc-70(cas983)* (N=62). Plotted as population sum ± 95% CI. Two-sided Fisher’s Exact Test **** p<0.0001, ns p=0.4671. (**B**) AFD-NRE microvilli length (μm) for WT (mean=12.23 ± 1.08 95% CI, N=13), *unc-23* (mean=16.91 ± 2.11 95% CI, N=12), *sma-1* D1 adults (mean=19.19 ± 3.46 95% CI, N=9) and *sma-1* D3 adults (mean=23.56 ± 6.73 95% CI, N=8). Plotted as mean ± 95% CI. Sidak’s Multiple Comparisons Test ** p=0.0091 (WT-*unc-23* D1), ** p=0.0040 (WT-*sma-1*D1), and ns p=0.1734 (*sma-1* D1-*sma-1* D3). (**C-E**) AMsh glial apical domain fluorescence labeling in WT (C) and *sma-1* mutants (D), quantified in (E). Apical domain (dotted magenta line) expands in mutants (arrow). Asterix = anterior pharyngeal bulb midpoint, # = glial soma. Scale bar, 8μm. (E) Quantification follows Figure 3P. Wild type (N=159), *unc-23* (N=92), *sma-1* (N=89). Two-sided Fisher’s Exact Test **** p<0.0001. (**F**) Mosaic rescue analyses. % AFD-NRE meander in *sma-1* mutants with unstable extrachromosomal arrays of wild type SMA-1 tracked with cell-specific co-injection markers. N≥88. Two-sided Fisher’s Exact Test ns p=0.7169, *** p=0.0002. (**G-J**) Schematic and fluorescence images of SMA-1:ZF1:GFP in WT (G, I), *zif-1(gk117)* mutant with AMsh glia-specific ZF1-degron (H), and *unc-23(e25)* mutant animals (J). Arrow = “mesh” staining in glia, lost in glia-specific ZF1 degron, and *unc-23*.blue # = unaffected SMA-1:GFP staining in epithelia. Scale bar, 8μm. (**K**) SMA-1 Mean Gray Value quantified in WT (1,272 ± 229 95% CI, N=23) and *unc-23* (226 ± 91 95% CI, N=10). Same day random sample imaging (100% T FITC, 0.20318s exposure). Two-tailed Welch’s t-test **** p<0.0001. (**L-N**). AMsh glial LifeAct reporter F-actin staining in WT (L) *unc-23* mutant (M) and *sma-1* mutant animals (N) shows aberrant staining in both mutants (pink hash). Scale bar, 8μm. Glial sheath (dotted purple).

Their overlapping AFD-NRE phenotypes led us to ask if UNC-23 and SMA-1 interact genetically by recombining *unc-23* with the *sma-1(ru18)* presumptive null allele (McKeown et al., 1998) (Figure S4B). *unc-23(e25) sma-1(ru18)* double mutant defects were indistinguishable from *unc-23(e25)* (Figure 4A), indicating that *sma-1* is genetically epistatic to *unc-*23. Since SMA-1 is large 482 kDa/4166 amino acid protein that may require Hsp chaperone activity to fold properly, we then wondered if it is a direct UNC-23 folding substrate. If so, we reasoned it must function in the same cell i.e. epithelia. To test this, we performed genetic mosaic cell-specific rescue experiments by coupling the SMA-1 rescuing fosmid with either epithelia or glia cell-specific marker. To our surprise, expression of *sma-1* bearing arrays in glia completely rescued the mutant AFD-NRE defects, while expression in epithelia did not (Figure 4F). Thus, while genetically epistatic, epithelial UNC-23 cannot directly regulate glial SMA-1 folding.

To confirm this, we examined SMA-1 protein expression with single-cell resolution *in vivo* using a single-copy *sma-1:zf1:gfp* transgene (Sanchez et al., 2021). In embryos, this transgene expresses in many tissues including epithelia in embryos (Praitis *et al*., 2005; Sanchez *et al*., 2021). Day 1 adults exhibit expression in restricted regions, including the pharyngeal, uterine spermatheca-uterine and rectal valves (Figure S4G-J) and possibly PHsh glia (S4K). In the anterior head region, staining was observed where hyp3/hyp4/hyp5 cells contact AMsh glia anterior regions around glia: AFD-NRE contact site zone (Figure 4G). Thus, SMA-1 is expressed in the appropriate anatomical region and age. The close packing of thin cytoplasmic processes of epithelia and glia at the animal’s nosetip preclude clear delineation of SMA-1:GFP at cell-specific resolution by fluorescence double-labeling (data not shown). Therefore, we utilized the ZF1/ZIF-1 degron system to aid in analysis of SMA-1:GFP expression (Armenti et al., 2014). Proteasomal degradation of SMA-1:ZF1:GFP in AMsh glia by cell-specific expression of *zif-1* revealed that the SMA-1 “mesh” staining was within AMsh glia (Figure 4H). Strikingly, this mesh was severely depleted in *unc-23* mutant animals (Figure 4I, 4J, 4K). Thus, epithelial UNC-23/BAG2 regulates glial SMA-1/β-Spectrin cytoskeleton *in vivo*.

Finally, since spectrins bind F-actin networks, we also examined distribution of F-actin in glia of *unc-23* mutant animals. For this, we engineered an AMsh-specific LifeAct F-actin reporter transgene (Spracklen et al., 2014). While wild-type animals express LifeAct in AMsh anterior domains, this staining was strikingly aberrant in both *unc-23* and *sma-1* mutant animals (*wild-type*, n= 10/11 animals; *unc-23*, n= 1/5 animals; *sma-1*, n= 1/10 animals) (Figure 4L-N). Thus, epithelial UNC-23 regulates AMsh glial cell shape by regulating its apical SMA-1/β-Spectrin and F-actin cytoskeleton.

### Epithelial UNC-23 regulates glial SMA-1 through EGL-15/FGFR

How does loss of epithelial UNC-23 function transmit to AMsh glia to regulate glial SMA-1? Given that UNC-23 is a co-chaperone with HSP-1, we first asked if this was triggered by general proteotoxic stress in *unc-23* mutant epithelia. However, administration of either of two proteostasis blocker drugs, MG-132 or Bortezomib/Velcade, did not recapitulate *unc-23* defects (Figure 5A). Epithelial expression of human α-synuclein; or epithelia-specific genetic block of proteosome function by expression of the mutant proteosome PBS-5^T65A^/PSMB (Lehrbach and Ruvkun, 2019) also did not induce AFD-NRE defects (Figure 5A). In corollary, there was no aberrant upregulation of ER or cellular stress response in *unc-23* mutants, as evaluated by the *hsp-4:gfp* or *P_rpt-3_:gfp* reporters, respectively (Figure S5A-S5B’’). In line with this, mutations that abrogate either ER stress response pathway (*pek-1, ire-1, xbp-1, atf-6)*, autophagy pathway (*unc-51, bec-1),* heat-shock response factor (*hsf-*1), or ubiquitin fusion degradation pathway (*cdc-48, ufd-2)*, did not phenocopy *unc-23(e25)* AFD-NRE defects (Figure S5C, S5D). Thus, we concluded that *unc-23* neural defects are likely not due to general cellular proteotoxic stress or upregulation of canonical cell stress-response pathways.

**Figure 5.**
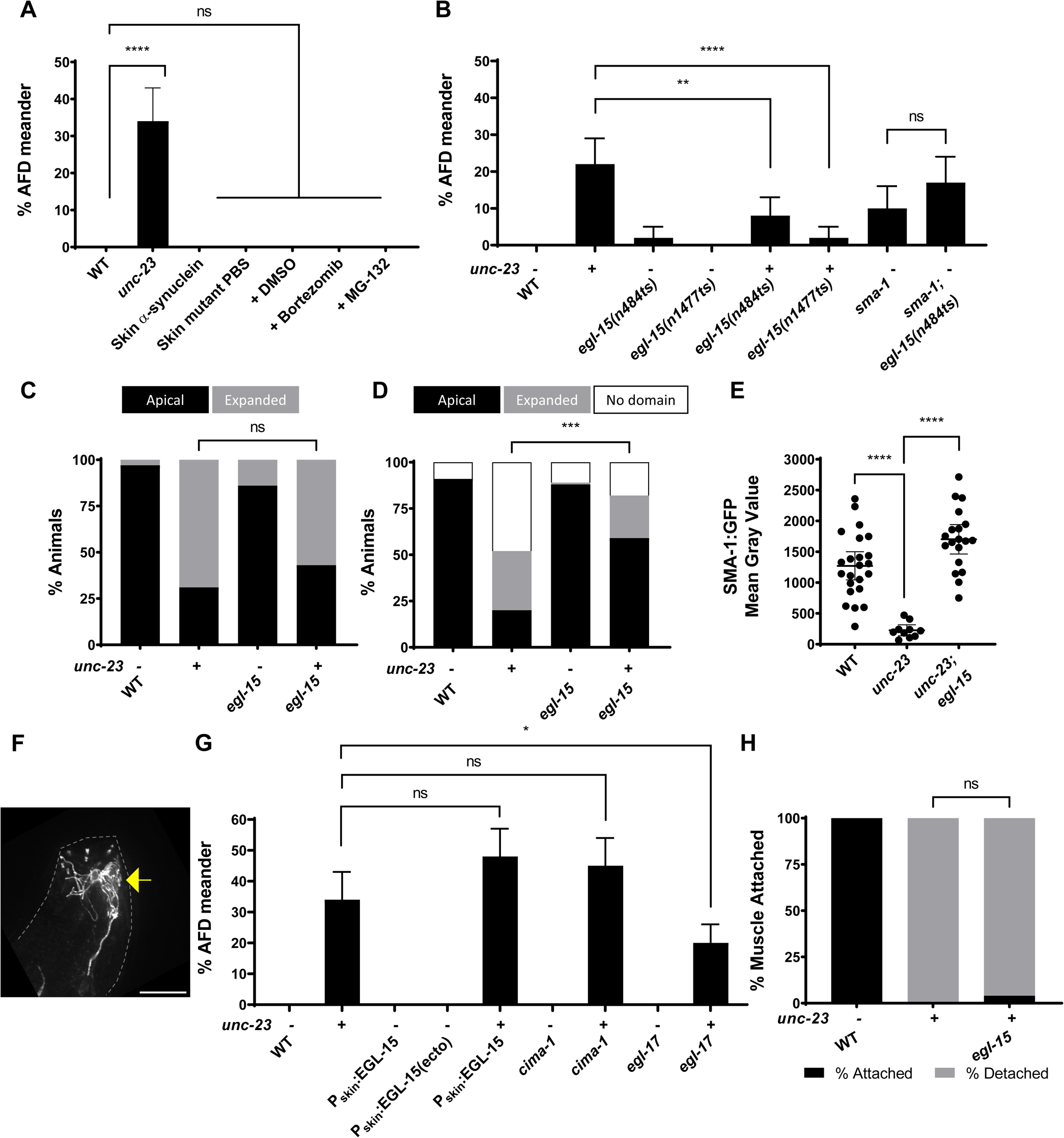
Epithelial UNC-23 regulates glial SMA-1 through EGL-15/FGFR signaling. Analyses on *unc-23(e25)*, *sma-1(ru18)*, and *egl-15(n484ts)* alleles unless noted, *nsIs645* AFD-NRE marker. (**A**) % AFD-NRE meander under different proteotoxic stress conditions, as noted. Wild type α-synuclein (N=35), PBS-5^T65A^ (N=56), MG-132 (N=56), Bortezomib (N=85). N≥112 for other genotypes. Plotted as population sum ± 95% CI. Two-sided Fisher’s Exact Test **** p<0.0001 and ns p>0.9999. (**B-D, G**). (B) % AFD-NRE meander microvilli, N≥100 for all genotypes except WT (N=81) and *egl-15(n484ts)* (N=97). Plotted as population sum ± 95% CI. Two-sided Fisher’s Exact Test ** p=0.0026, **** p<0.0001, and ns p=0.1625. (C) Quantification of skin epithelia apical enrichment. Wild type (N=68), *unc-23 (*N=49), *egl-15* (N=50), *unc-23; egl-15* (N=95). Two-sided Fisher’s Exact Test ns p=0.2851. (D) Quantification of AMsh apical enrichment. WT (N=106), *unc-23* (N=107), *egl-15* (N=92), *unc-23; egl-15* (N=118). Two-sided Fisher’s Exact Test *** p=0.006. (**E**) Quantification of SMA-1 Mean Gray Value in WT (1,272 ± 229 95% CI, N=23), *unc-23* (226 ± 91 95% CI, N=10) and *unc-23; egl-15* (1,703 ± 240 95% CI, N=19). Dunnett’s T3 Multiple Comparisons Test **** p<0.0001. (F) Representative image of SMA-1:GFP glial web (yellow arrow) restored in *unc-23; egl-15* mutants. Scale bar, 8μm. (G) % AFD-NRE meander microvilli with epithelia-specific EGL-15^A^ (N=95), EGL-15A^ecto domain^ (N=98), EGL-15^A^ in *unc-23* (N=101). *cima-1(wy84)* and *egl-17*(*ay6)* alleles used. N≥112 for all other genotypes except *egl-17* (N=72). Data plotted as population sum ± 95% CI. Two-sided Fisher’s Exact Test ns p=0.0553 [(*unc-23(e25)*-P*_dpy-7_*:EGL-15; *unc-23(e25)*], ns p=0.0934 [*unc-23(e25)* - *cima-1(wy84)*; *unc-23(e25)*], and * p=0.0104 [*unc-23(e25)* - *egl-17(ay6)*; *unc-23(e25)*]. (H) % anterior muscle attachment in WT (N= 27), *unc-23* (N= 10), and *unc-23; egl-15(n1477ts)* (N=27). Two-sided Fisher’s Exact Test ns p>0.9999.

To define how epithelia:glia communicate to maintain glial polarity and neuron shape, we next performed a candidate screen of known signaling factors. Mutations in dense core vesicle/neuropeptide signaling (*unc-31* CADPS or *egl-21*/carboxypeptidase E) (Figure S5E), serine convertases (*kpc-1, bli-4)* neither phenocopied nor suppressed *unc-23* mutant AFD-NRE defects (Figure S5E-S5G). Mutations in major neurodevelopmental pathways are also dispensable for AFD-NRE shape control (*unc-40/*DCC, *sax-7/*L1CAM, *unc-6*/netrin, *unc-129*/TGFβ, *unc-71*/ADAM10 metalloprotease, *smp-1/smp-2/*semaphorins) (Figure S5H-S5K).

Strikingly, loss-of function mutations in EGL-15/FGFR strongly suppressed *unc-23* mutant AFD-NRE defects (Figure 5B). The *egl-15* gene locus produces alternatively spliced transcripts, *egl-15^A^* and *egl-15^B^*, with EGL-15^A^ predominant in neural roles (Figure S5L) (Bülow et al., 2004; Burdine et al., 1997; Goodman et al., 2003; Shao et al., 2013). A mutation that depletes only EGL-15^A^ [*egl-15(n484)*], suppressed AFD-NRE defects as well as one that affects both transcripts [*egl-15(n1477)*] (Figure 5B). Thus, EGL-15^A^ mediates UNC-23 function.

To place EGL-15^A^ gene function in the tri-cellular communication cascade, we asked if it can similarly suppress *sma-1* mutant defects and found that it did not (Figure 5B). This genetically placed EGL-15^A^ function downstream of epithelial UNC-23 but upstream of glial SMA-1 functions. Prior studies have shown that EGL-15/FGFR function in epithelia provides substrate for cell spreading and axon outgrowth (Bülow *et al*., 2004; Shao *et al*., 2013). So, we hypothesized that EGL-15^A^ may enable cellular stretching of either skin epithelia and/or glia. To test this, we examined skin epithelia and glial apical domains in *unc-23* animals with and without *egl-15^A^* lesion. While skin epithelia apical domain expansion of *unc-23* mutant animals is unaffected by loss of *egl-15,* glial apical domain defects were significantly rescued (Figure 5C, D). Thus, similar to AFD-NRE defects, EGL-15 functions downstream of epithelial apical domain expansion/stretching, and upstream of glial apical domain defects. Lastly, we examined how EGL-15 impacts SMA-1 levels. Loss of SMA-1: GFP levels in *unc-23* mutant animals was strikingly restored in *unc-23; egl-15* double mutants (Figure 5E, 5F). Thus, EGL-15^A^ relays *unc-23* epithelia stretching to regulate glial SMA-1 and apical domain expansion, and consequently informs AFD-NRE shape.

Is EGL-15 permissive or instructive in inducing glial apical domain expansion? Over-expression of either EGL-15^A^ or EGL-15^A^ ectodomain in epithelia, or mutation in the epithelial EGL-15 negative regulator CIMA-1 (Shao *et al*., 2013) did not impact AFD-NRE, suggesting a different cellular mechanism is at play (Figure 5G). Thus, in animals with normally delimited skin epithelia apical domains, increasing levels of EGL-15 signaling alone cannot drive glial cell-shape defects. However, in animals with aberrant epithelial apical polarity such as in *unc-23*, it permissively enables glia-neuron overgrowth.

Finally, EGL-15/FGFR can have both ligand-dependent and non-canonical mechanisms of action (Bülow *et al*., 2004; Goodman *et al*., 2003). Mutations in the *egl-17/FGF* ligand also suppressed *unc-23* AFD-NRE defects (Figure 5G), suggesting that canonical ligand binding drives this function. Interestingly, *egl-15* did not suppress muscle detachment defects of *unc-23 (e25)* (Figure 5H). Thus, EGL-15/FGFR and EGL-17/FGF are molecular transducers of specifically epithelia-to-glia/neuron defects in *unc-23* mutants.

### Blocking glial secretome or neuron activity mitigates UNC-23 AFD-NRE defects

Having dissected how epithelia communicate to glia, we next asked if AMsh glia communicate to AFD-NRE to causally induce its overgrowth in *unc-23* mutant animals. Ablation of the glial cell via cell-specific diptheria toxin expression showed attenuation of AFD-NRE overgrowth (data not shown) (Bacaj *et al*., 2008b; Singhvi *et al*., 2016), as did knockdown of the glial secretome regulator PROS-1/Prox (Figure 6A) (Kage-Nakadai et al., 2016; Wallace *et al*., 2016). These results are consistent with glia mediating UNC-23’s regulation of AFD-NRE shape.

**Figure 6.**
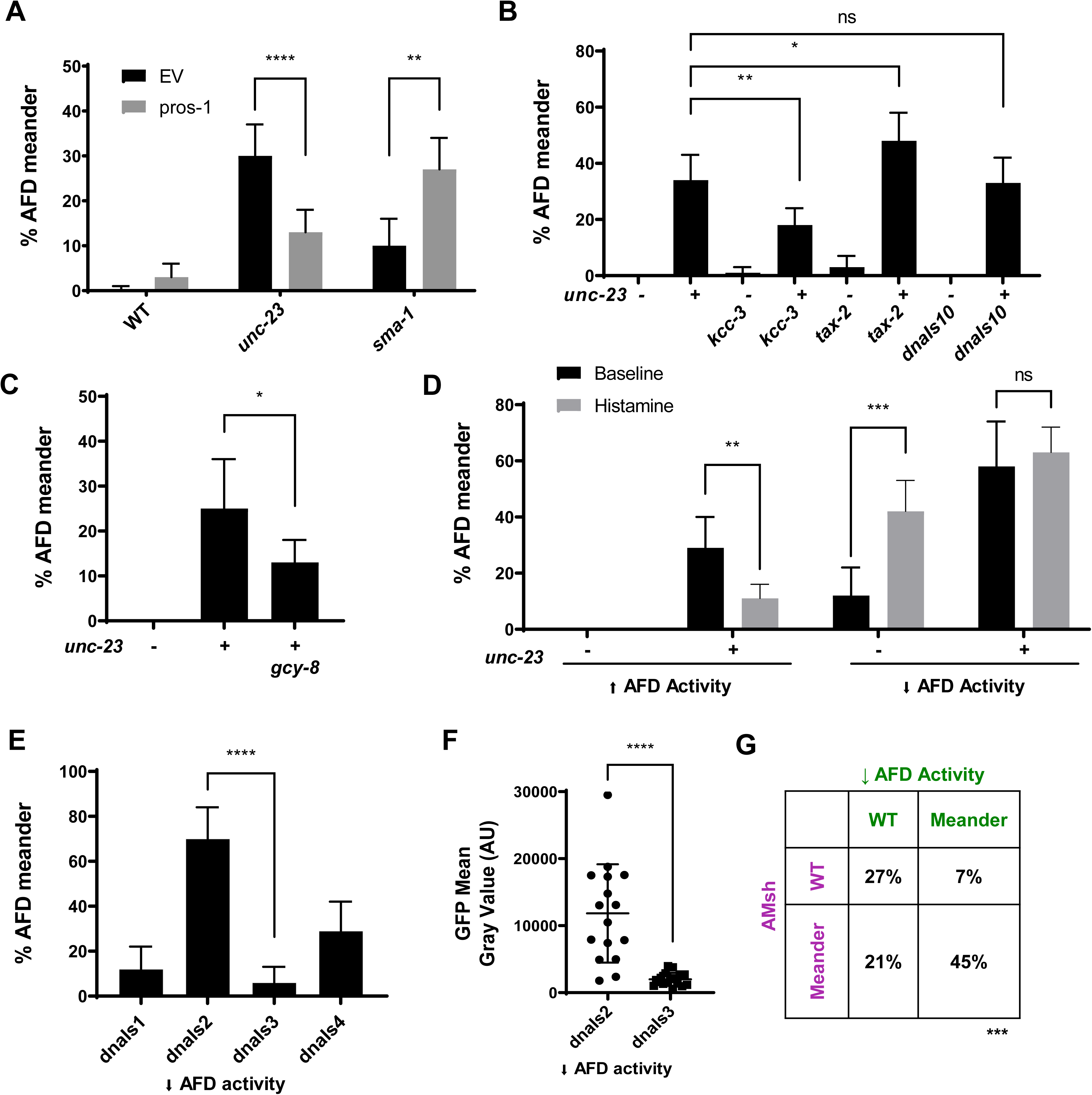
Modulating glial secretion or neuron activity can bypass *unc-23* neural defects. (**A**) % AFD-NRE meander in WT, *unc-23(e25)*, and *sma-1(ru18)* mutants in control or *pros-1* RNAi visualized by *nsIs645*. Plotted as proportional sum of three independent RNAi rounds ± 95% CI. N≥135 for all conditions and genotypes except *sma-1* mutants on empty vector control (N=94). Two-sided Fisher’s Exact Test **** p<0.0001 and ** p=0.0013. (**B**) % AFD-NRE microvilli meander in *unc-23(e25)*, *kcc-3(ok228), tax-2(p691), and dnaIs10 [P_AMsh_:KCC-3]*. N≥92 for all genotypes except *tax-2* (N=69) and *dnaIs10* (N=17). Plotted as population sum ± 95% CI. Two-sided Fisher’s Exact Test ** p=0.0027, * p=0.0478, and ns p=0.8850. (**C**) % AFD-NRE microvilli meander in *unc-23(e25)*, *gcy-8(ns335)* and *unc-23(e25); gcy-8(ns335)* visualized with *nsIs228*. N≥67 for all genotypes. Plotted as population sum ± 95% CI. Two-sided Fisher’s Exact Test * p=0.0277. (**D**) Chemo-genetic manipulation of AFD neuron activity using histamine-gated cation or chloride channels in WT and *unc-23(e25)*. N≥68 for all conditions except WT decreased activity baseline (N=40), WT increased activity base line (N=41), and *unc-23* decreased activity baseline (N=33). Plotted as population sum ± 95% CI. Two-sided Fisher’s Exact Test ** p=0.0015, *** p=0.0009, and ns p=0.6836. (**E**) % AFD-NRE meander at baseline in four independent chemo-genetic silencing transgenic strains: *dnaIs1*(N=41), *dnaIs2* (N=44), *dnaIs3 (*N=35), and *dnaIs4* (N=51), plotted as population sum ± 95% CI. Two-sided Fisher’s Exact Test **** p<0.0001. (**F**) Quantification of AFD GFP mean gray value in *dnaIs2* (11,829 ± 3,907 95% CI, N=16) and *dnaIs3* (1,989 ± 498 95% CI, N=17) from same-day random sample imaging (50%T FITC, 0.005s exposure). Data is mean ± 95% CI. Welch’s t-test **** p<0.0001. (**G**) Correlation of *dnaIs2* AFD low activity meanders with AMsh glial meander (N=73). Two-sided Fisher’s Exact Test *** p=0.0001.

We and others have previously shown that AMsh glia regulate AFD-NRE by multiple independent mechanisms, including pruning and through glial transporter KCC-3/SLC12A6, (Raiders *et al*., 2021a; Raiders et al., 2022; Singhvi *et al*., 2016; Yoshida et al., 2016). We therefore wondered if glia transduce AFD-NRE overgrowth in *unc-23* mutant animals through either defective pruning or altered glial KCC-driven signaling. We found no change in steady-state number of pruned AFD-NRE fragments in *unc-23* mutant animals, suggesting this was likely not the driver (SAR, personal communication.). *kcc-3* mutants, however, exhibit AFD-NRE collapse defects and suppressed *unc-23(e25)* meander defects (Figure 6B). Further, over-expression of KCC-3 does not phenocopy *unc-23* or exacerbate its effects (Figure 6B). *kcc-3* mutants have elevated second messenger cGMP in AFD (Singhvi *et al*., 2016). So, we next asked if the observed genetic suppression reflects their role in AFD neuron’s cGMP modulation. Indeed, mutations in a gain-of-function allele of *gcy-8 (ns335)* (increased neuronal cGMP) also suppress *unc-23* defects (Figure 6C). Taken together, we infer that cGMP levels in AFD-NRE, not levels of glial KCC-3 around AFD-NRE, mechanistically link KCC-3 and UNC-23 in driving AFD-NRE.

cGMP in the AFD-NRE regulates AFD activity through cyclic nucleotide-gated channels (Cho et al., 2004; Inada *et al*., 2006; Satterlee et al., 2004; Singhvi *et al*., 2016; Yoshida *et al*., 2016). This led us to wonder if suppression by altered cGMP levels was due to its modulation of AFD neuron activity. To test this, we perturbed AFD activity independent of cGMP in three ways. One, by examining mutations mutations in TAX-2/CNG channel β-subunit (reduced neuron activity). We found that these alone have slightly mis-shapen AFD-NRE and only mildly enhance *unc-23* AFD-NRE defects (Figure 6B) (Komatsu et al., 1996; Satterlee *et al*., 2004; Singhvi *et al*., 2016). Two, by chemo-genetic activation of AFD in adulthood. For this, we engineered transgenic animals that enable manipulation of AFD neuron activity upon exogenous dietary supplementation of Histamine using histamine-gated ion channels (Pokala et al., 2014). We found that neuron hyper-activation using a Histamine-gated cation channel suppressed *unc-23* AFD-NRE defects (Figure 6D). Three, by chemo-genetically silencing of AFD using histamine gated chloride channels (Pokala *et al*., 2014). We found that this alone resulted in AFD-NRE meander defects reminiscent of *unc-23* loss (Figure 6D), with defect severity strikingly tracking transgene expression levels as measured by fluorescence intensity of a bi-cistronic GFP reporter (Figure 6E, 6F). Strikingly, this does not enhance *unc-23* mutant defects. For these chemo-genetic studies, we induced inactivation at the last larval L4 molt, and note that this was sufficient to alter AFD-NRE shape of adult animals. This is parallel corroboration that L4-adult molt development critical period (and not only embryonic development) regulates AFD-NRE shape in adults. Finally, to ask if reduced AFD activity can reciprocally alter glia-shape, we examined AMsh glia by fluorescence labeling in chemo-genetically inactivated AFD transgenic animals. Meandering AFD-NRE in these animals also exhibited meandering of AMsh glia (Figure 6G) but not muscle detachment (data not shown). Thus, AFD’s cGMP levels and thereby activity regulate AFD-NRE shape downstream of UNC-23/BAG2, and activity alone can reciprocally influence AMsh glial cell shape.

### UNC-23 reveals heterogeneity in epithelia-glia interactions

Anterior head epithelial cells contact multiple glia-neuron pairs and AMsh glia contact other NREs (Figure 7A). We therefore asked if the UNC-23/EGL-15/SMA-1 pathway also regulates other glia-neuron units. Cell-specific reporter analyses reveal that NREs of all neurons embedded within AMsh glia (AWA, AWB, AWC) exhibit analogous “meander” defects (Figure 7A-E). That other AMsh-embedded NREs meander may also explain why we found that AMsh glia meander even if AFD-NRE do not, they may be tracking other NREs (Figure 1M, 7A). In contrast, shape of an NREs that traverses the AMsh glia: AMso glia channel (ASE-NRE) is unaffected by *unc-23(e25)* lesion (Figure 7A, D-D’, E) (Perkins *et al*., 1986; Ward *et al*., 1975)

**Figure 7.**
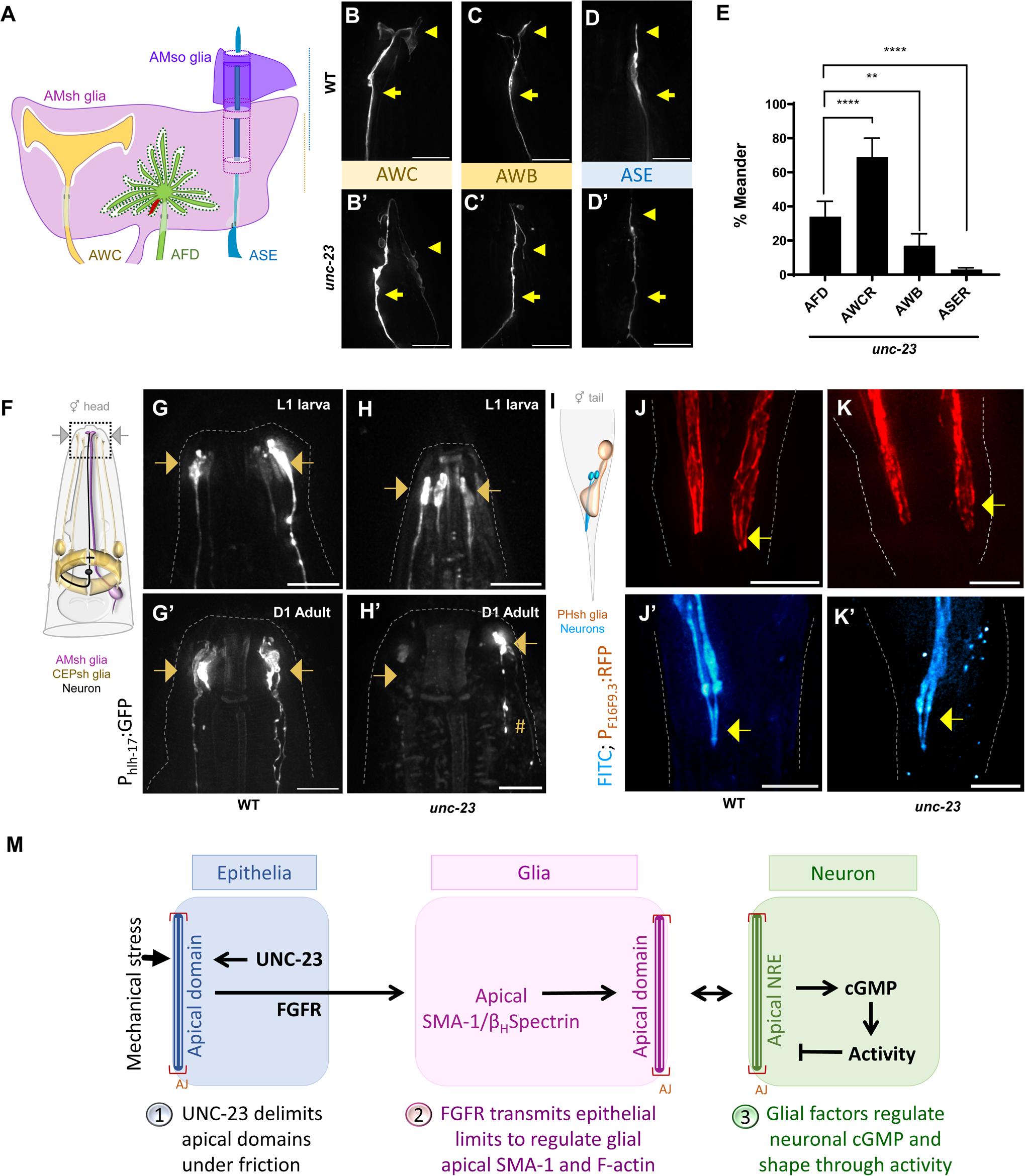
UNC-23 reveals regional heterogeneity in epithelia-glia-neuron cross-talk. (**A**) Schematic of AMsh glial contact with AMso glia, embedded neurons (AWC) and channel neurons (ASE). (**B-D’**) Fluorescence images of AWCR (B, B’), AWB (C, C’), and ASER (D, D’) in WT (B-D) and *unc-23* (B’-D’). Scale bar, 8μm. Dendrites (arrow) and NREs (arrowheads) denoted (**E**) % *unc-23(e25)* meander in AFD (N=122), AWCR (N=61), AWB (N=106), and ASER (N=70) neurons. Plotted as population sum ± 95% CI. Two-sided Fisher’s Exact Test **** p<0.0001 and ** p=0.0040. (**F-H’**) Schematic (F) of animal head, boxed region imaged in G-H’. Fluorescence images of boxed region in F. (G-H’) of CEPsh anterior endings in L1 larva (G, H) and adult worms (G’, H’, box 2 in F) in WT (G, G’) and *unc-23* mutants (H, H’). Scale bar, 5μm. Dim/missing CEPsh processes (arrows) and meandering apical endings (arrow) noted. (**I-K’**) Schematic (I) and fluorescence images (J-K’) of phasmid neurons (J, J’) and PHsh glia (K, K’) in the tail of the animal in WT (J, K) and *unc-23* mutants (J’, K’). Phasmid neurons stained by FITC. Scale bar, 5μm. Phasmid sheath glia (red) process (arrow) with phasmid neurons (cyan) are normal in *unc-23(e25)* mutants. Scale bar, 5μm. (**L**) Model. Epithelia impact glia-neuron shape under mechanical stress via the UNC-23/FGFR/Spectrin molecular pathway.

We also examined glia in two other sense-organ pairs, the anterior head-region located CEPsh glia and posterior tail-located PHsh glia. CEPsh glia extend anterior projections to the animals’ nose-tip posterior sheaths that wrap and infiltrate the nerve ring (Figure 7F-H) (Singhvi and Shaham, 2019; White et al., 1986). We found variable and pleiotropic defects in every CEPsh glia examined (n=22/22), including dim/missing cells and processes, meandering CEPsh anterior endings, and CEPsh process detachment (Figure 7G’-H’). We also noted that CEPsh posterior processes, which envelop the animal’s “brain” neuropil, extended aberrant posterior projections in *unc-23* mutant animals (Figure S6). Intriguingly, while CEPsh glia defects in mutant animals, like AMsh glia defects, are age-dependent (normal CEPsh in L1 larvae, n=10/10), defects in the two glia are not phenotypically identical (Figure 7G-H’ versus Figure 1I-K).

Surprisingly, neither the posteriorly placed phasmid neurons nor their contacting PHsh glia had any shape defects (Figure 7I-K’). This contrast yields two insights. One, there is heterogeneity in epithelia-glia interactions, such that distinct glia in the same anatomical region are differently regulated by epithelial UNC-23/BAG2 (anteriorly placed AMsh vs CEPsh). Two, epithelial UNC-23 control of glia-neuron shape exhibits regional bias across the nervous system (anteriorly placed AMsh and CEPsh vs posteriorly placed PHsh), despite PHsh glia expressing the downstream SMA-1/β Spectrin (Figure S4J).

## DISCUSSION

How glial-neuron units maintain their shape life-long is unknown. This study finds that epithelia maintain glial cell-polarity and glia-neuron shapes against mechanical shear, and uncovers the underlying molecular mechanism (Figure 7L). Mechanistically, loss of epithelial UNC-23/BAG2 causes aberrant expansion of epithelial apical domains under mechanical shear from animal locomotion. Through FGFR, this causes loss of apical β-Spectrin and F-actin cytoskeleton in contacting glia, causing glial shape and polarity defects. In turn, through secreted glial factors, this causes aberrant overgrowth of some associated NREs that appose glial apical membranes. Impairing mechanical strain, glial secretion, or altering neuron activity, mitigates *unc-23* mutant animal neural defects. Furthermore, this study finds that epithelial regulates distinct glia-neuron pairs differently, revealing both regional specificity and cellular heterogeneity in epithelia-glia interactions. Lastly, epithelial UNC-23 acts within a developmental critical window to maintain glia-neuron and sense-organ shape lifelong. Conservation of all molecular components identified here, and analogy of epithelia-glia-neuron anatomical orientation, lead us to suggest that similar mechanisms may broadly govern epithelial modulation of glia-neuron shape maintenance, with impact on aging and disease.

### Analogous epithelia-glia-neuron contacts across systems and species

Our work shows that epithelia communicate with glia to dictate glial cell-polarity and shape, with consequence for neuron shape and activity; and uncovers the underlying tri-cellular molecular cascade. We note that epithelia-glia-neuron contacts bear striking analogy to epithelia:AMsh:AFD in their cellular orientation across systems and species, including sense-organs, blood-nerve/brain barrier, and glia-meningeal contacts. In the instances examined, apical membranes of glia contact neurons and glial basolateral domains (or markers thereof) enrich at epi/endothelia or basement membrane contact-sites (Derouiche *et al*., 2012; Low *et al*., 2019; May-Simera and Kelley, 2012; Salzer, 2003; Silies and Klambt, 2011). Thus, *Drosophila* ensheathing glia apical membranes contact neurons while basolateral membranes face the blood-brain barrier (Pogodalla et al., 2021). This polarity is regulated by glial spectrin, loss of which alters animal behavior (Pogodalla *et al*., 2021). Vertebrate retinal Müller glia apical domains contact photoreceptors, and basolateral end-feet contact blood vessel endothelia (Derouiche *et al*., 2012; Vecino et al., 2016). Mammalian Schwann cell glia orient apical membranes to neurons and basolateral domains to basal lamina (Derouiche *et al*., 2012; Etienne-Manneville, 2008; Salzer, 2003; Tricaud, 2017). Mammalian CNS astrocyte end-feet localize basolateral markers like AQP4 to the blood-brain barrier, while their processes contact neurons (Derouiche *et al*., 2012; Etienne-Manneville, 2008; Tricaud, 2017). Lastly, radial glia and astrocytes contact pial surfaces basolaterally, while radial glial apical end-feet contact ventricles (Derouiche and Geiger, 2019; Taverna *et al*., 2014). Functionally, *C. elegans* epithelia dictate glia-dependent synapse positioning, although cell-polarity has not been examined in this model (Shao *et al*., 2013). Despite this parallel, how heterotypic epithelia-glia-neuron cells coordinate their cell-organization, of the functional relevance thereof, is barely explored. Further, while there is evidence of glia-to-endothelia signaling (Derk *et al*., 2021; Kaplan et al., 2020; Langen *et al*., 2019), whether or how epi/endothelia signal to glia is less-described.

### Mechanical forces in neural function and disease

We find that this epithelia-glia-neuron signaling couples glia-neuron shape maintenance to mechanical forces sensed by the organ/animal. The following lines of evidence lead us to speculate that similar hetero-cellular mechanobiological coupling may also occur in both PNS and CNS. In the PNS, glia-neuron units embedded either in sensory epithelia (olfactory, gustatory, auditory) or skin epithelia (touch), that face external mechanical stress life-long. Interestingly, mutations in genes regulating cellular tensional homeostasis drive sensory dysfunction across sensory modalities (Lumpkin et al., 2010; Nolano et al., 2008; Phuong Le *et al*., 2022; Tarchini et al., 2016; Yadav et al., 2019). Mechanical stress on gut and bladder epithelia also modulates sensory efferent nerve activity (Najjar et al., 2020), and chemosensory neuron activity in *C. elegans* (Caprini et al., 2021), although glial coupling is not yet explored for either setting. Mechanical stress also drives sensory epithelium neurogenesis and placode development (Levy Nogueira et al., 2015; Phuong Le *et al*., 2022; Sicard et al., 1998). In the CNS, blood-flow mechanical shear impacts the blood-brain barrier, which is maintained by endothelia-glia interactions (Bauer et al., 2014; Bundgaard and Abbott, 2008; Segarra et al., 2018). Indeed, vascular hypertensive patients co-present blood-brain barrier dysfunction and astrogliosis (Abbott, 2002; Hasan-Olive et al., 2019) and intracranial or vascular mechanical stress/injury is linked to neurodegeneration. However, how mechanical tension is transmitted to neurons either directly or through glia, is poorly studied, and these correlative clinical findings await inquiry into molecular causality (Levy Nogueira *et al*., 2015).

The molecular machinery of epithelia-glia mechanobiology coupling we found is distinct from how *C. elegans* epithelia protect axon degeneration (no glia involved) or glia-dependent synaptogenesis (no mechanical input) (Bonacossa-Pereira et al., 2022; Coakley et al., 2020; Shao *et al*., 2013). Interestingly, components of this pathway (BAG2, FGFR, Spectrins) also regulate cell polarity and cancer epithelial-mesenchymal transitions (Antony et al., 2019; Lu and Lu, 2021; Perez-Moreno et al., 2003). Their levels track aggressiveness of cancers like glioblastoma (Behl, 2016; Gentilella and Khalili, 2011; Qin et al., 2021) that have abnormal vascularization and ECM mechanical properties (Khoonkari et al., 2022; Sun and Kim, 2022). This raises the possibility that this molecular cascade may broadly transduce epithelia-glia mechanobiological coupling, suggesting that its role in epithelia-glia signaling in sensorineural health, aging and disease warrants further inquiry.

### Cellular and temporal heterogeneity in epithelia-glia interactions

Besides uncovering epithelia-glia communication, our work also demonstrates previously unappreciated spatial and temporal heterogeneity in their interactions. This extends the emerging concept of glia heterogeneity from glia-neuron to also epithelia-glia interactions (Ben Haim and Rowitch, 2017; Zhang and Barres, 2010). Temporally, epithelia regulate glia-neuron shapes only in adult, through UNC-23 function at a critical developmental period, despite the anatomical organization being already established in the embryo. Spatially, our preliminary studies (data not shown), hint at three anterior epithelial cells as the presumptive site of action for AMsh, of which one (hyp4) gap-junctions with AMsh (Altun et al., 2009; Altun *et al*., 2002). Prior EM and fixed-tissue staining also reported spatio-temporal variance in epithelial cytoskeleton (Costa et al., 1997). Thus, it may be that the anterior bias of UNC-23 effects on glia (head AMsh and CEPsh vs tail PHsh) may derive from its specific roles in anterior epithelia. The variability of epithelial UNC-23’s impact on different anterior glia (AMsh vs CEPsh), however, remains puzzling. Whether this arises from divergent epithelial properties and/or different glial responsivity profiles, or even associated neuron activity states, will be an exciting avenue of future inquiry.

## ACKNOWLEDGEMENTS

We thank the following for generously sharing reagents: Ann Wehman (*pat-3* RNAi clone); Chun-Liang Pan (LifeAct reporter); Jessica Feldman and Lauren Cote (*sma-1* reporters); Nic Lehrbach (*P_col-10_:* PBS-5^T65A^); Navin Pokala and Cori Bargmann (chemogenetic His^Cl^ and His^Cat^ plasmids). We thank the Singhvi lab and Jihong Bai for discussions and comments on the manuscript; lab members Erik Black, Sneha Ray, and Maria Purice for gift of reagents; Sneha Ray and Stephan Raiders for sharing preliminary data as personal communication; and Maria Purice for guidance on RT PCR experiments. This work was funded by Simons Foundation/SFARI grant (488574), Esther A. & Joseph Klingenstein Fund and the Simons Foundation Award in Neuroscience (488574) and NIH/NINDS funding (NS114222) to AS. This work was performed while AS was a Glenn Foundation for Medical Research and AFAR Junior Faculty Grant Awardee. AS sincerely thanks philanthropic supporters to her laboratory. This project was initiated while AS was in Shai Shaham lab. Some work was performed at the Fred Hutch Shared Resources Core Facilities. We sincerely apologize if we missed citing works due to our oversight or space considerations.

## AUTHOR CONTRIBUTIONS

CM and AS designed, performed, and analyzed all experiments, along with JB. AS co-wrote the manuscript with CM, along with JB.

