## Supplemental Materials and Figures for "Epithelia delimits glial apical polarity against mechanical shear to maintain glia-neuron - architecture"

**This file contains:**

Experimental Methods and References

Supplemental Figures and Legends

### EXPERIMENTAL PROCEDURES

#### *C. elegans* methods

*C. elegans* were cultured as previously described(Brenner, 1974; Stiernagle, 2006). Bristol N2 strain was used as wild type. Animals were raised at 20ºC (unless noted) for two weeks without starvation. Temperature-sensitive *egl-15(n484)* and *egl-15(n1477)* alleles and relevant controls were grown at 20°C and shifted to 25°C at L4 (Bülow et al., 2004; Burdine et al., 1997). For analyses with *hsp-1(ra807)*, animals including relevant controls were grown at 15°C and shifted to 20°C at L4 (Rahmani et al., 2015). L4 larval animals were picked to fresh plates and assayed 24 hours later, unless otherwise noted. Germ-line transformations by micro-injection to generate unstable extra-chromosomal array transgenes were carried out using standard protocols (Mello et al., 1991). Integration of extra-chromosomal arrays was performed using UV with or without trioxalen (Sigma, T6137). All transgenic arrays were generated with 5 ng/µl $P_{elt\text{-}2}$:mCherry, 30 ng/µl $P_{mig\text{-}24}$:Venus, or 20 ng/µL $P_{unc\text{-}122}$:RFP as co-injection markers (Abraham et al., 2007; Armenti et al., 2014; Miyabayashi et al., 1999). Information on all strains and reagents is available by request. Dye-filling was performed per standard protocols (Hedgecock et al., 1985).

#### Strains

Strains were sourced from (a) the CGC, funded by NIH Office of Research Infrastructure Programs (P40 OD010440), (b) the International C. elegans Gene Knockout Consortium (C. elegans Gene Knockout Facility at the Oklahoma Medical Research Foundation, funded by the National Institutes of Health; and the C. elegans Reverse Genetics Core Facility at the University of British Columbia, funded by the Canadian Institute for Health Research, Genome Canada, Genome BC, the Michael Smith Foundation, and the National Institutes of Health) and (c)

National BioResource Project (NBRP), Japan.

**Mutagenesis, mapping of genetic lesions, protein structure predictions**

Mutagenesis and mapping protocols were as previously described (Shaham, 2007; Singhvi et al., 2016; Wicks et al., 2001). Briefly, *nsIs228* ($P_{srtx\text{-}1}$:GFP) animals were mutagenized with 75mM ethyl methane sulfonate (EMS, Sigma M0880) for 4 hours at 20ºC. 10,800 non clonal F2 progeny were screened for defects in AFD-NRE shape on a Zeiss Axioplan2 fluorescence microscope with 63x/1.4 NA objective and dual-band filter set (Chroma, Set 51019). Mutant animals were recovered from the slide were cloned. *unc-23(ns344)* was identified by a combination of SNIP-SNP mapping to a 0.63cM interval (Wicks *et al.*, 2001), whole genome sequencing, fosmid rescue and candidate gene analyses. Recessivity was determined by cloning individual animals and assaying neuronal phenotype segregation in F1 progeny. Briefly, 33/33 animals with meandering AFD-NRE segregated Unc progeny with either collapsed, meandering, or wild type AFD-NRE. In contrast, 4/21 animals with collapsed AFD-NRE segregated non-Unc progeny and 10/21 segregated progeny with AFD-NRE defects. Protein structures were predicted using Intensive mode Phyre2 (Kelley et al., 2015).

**Plasmids**

UNC-23**:** cDNA for UNC-23A/B/C isoforms were PCR amplified from a mixed stage cDNA library and cloned into pAS423 using Xma1/Nhe1 sites to generate pAS426. Subsequent manipulations were cloned via PCR amplification of unique promoter regions, or site-directed mutagenesis of cDNA.

$P_{glia}$: or $P_{epithelia}$:ApiGreen (apical membrane markers): Truncated SAX-7: sfGFP sequence from pMH510 (Low et al., 2019) was inserted into pAS465 and ASJ15/pCM1 using SphI/BamHI enzymes to generate ASJ74/pCM7 and ASJ76/pCM9.

$P_{epithelia}$:α-Synuclein: WT α-Synuclein sequence was codon optimized for *C. elegans* (Redemann et al., 2011) and synthesized by Biomatik as PB4147K. PB4147K was inserted into ASJ15/pCM1 using Xma/EcoRI to generate ASJ128/pCM17.

$P_{AMsh}$:LifeAct: 0.6kb sequence 5' of the F53F4.13 predicted coding region was PCR amplified from ASJ70/pEB009 with flanking Xba/Xma digestion sites and inserted into LifeAct:mNeonGreen (Huang et al., 2020) to generate ASJ95/pEB011.

$P_{AMsh}$: or $P_{epithelia}$:worm-mScarlet: Codon-optimized worm mScarlet was synthesized by Genewiz with flanking AgeI/EcoRI digestion sites and engineered into pAS545/pAB44 in a two-step process to create ASJ39/pMP6. To produce ASJ223/pCM25, ASJ15/pCM1 was restriction digest-cloned into ASJ39/pMP6 with BamHI/SpeI.

$P_{AFD}$:Histamine-gated cation channel (His-Cat): pNP497 (gift from N. Pokala/C. Bargmann) was digested with NheI/KpnI restriction enzymes to isolate the histamine-gated cation channel sequence and ligated to pAS178 to produce pAS537.

**RT-PCR**

RNA for three independent biological replicates was isolated following TRIzol LS and 5PRIME

Heavy Phase Lock Gel protocols, with modifications. Briefly, worms in TRIzol were vortexed at RT every 2 minutes for 15 minutes and placed at -80°C before RNA extraction. RNA was purified via Qiagen RNeasy Mini elute kit, quantified via nanodrop, normalized to 100ng, and cDNA was obtained using QuantiTect RT kit. RT PCR was performed in two technical replicates on WT and *ok1408* mutant cDNA without dilution. A forward primer within the *ok1408* deletion (5' CTCTCAGCAGAATCAAGCTCCTCC) and a forward primer at the beginning of exon 5 (5' GCCTACAAACCTACATGAACGCC) were separately paired with and a reverse primer at the end of exon 5 (5' GGATCCCTATTCGCTTTGATCATCC).

**Statistical Methods**

All data and statistics were graphed and analyzed using Prism9. Proportional data is presented as the proportional sum across multiple days of data collection ± 95% confidence interval. Two-Sided Fisher's Exact tests were performed to compare across genotypes, drug conditions, or RNAi treatments. For microvilli/cilia length measurements and fluorescence intensity measurements, Grubbs and Shapiro-Wilk test were used to identify outliers and test for normality. Brown Forsyth and Welch ANOVA with Dunnett's T3 multiple comparisons test (Microvilli) or Welch's unpaired t-test (cilia/ Fluorescence Intensity) were used to compare across genotypes. Kaplan Meyer curves and Lifespan Statistics were generated using Oasis2 (Han et al., 2016) via a Log Rank (Mantel-Cox) test.

**Microscopy, Image Processing and Analysis**

Worms were immobilized with 40mM sodium azide. Images were collected on a Deltavision Elite RoHS wide-field deconvolution system with Ultimate Focus (GE), a PlanApo 60×/1.42 NA or OLY 100×/1.40 NA oil-immersion objective and a DV Elite CMOS Camera. Some images were also collected on a Leica SP8 DLS inverted confocal microscope with a 100x/1.4 HC PL APO STED oil-immersion objective and HyD SMD 4/PMT Trans detector. Image processing was done in FIJI ImageJ. All background subtractions (50-pixel unless otherwise noted) were performed on both WT and mutant for consistency. Images for fluorescence intensity measurements were taken on the same day using random sampling. His-Chloride images were taken at 50% T with a 0.005s exposure and SMA-1:GFP images were taken at 100% T with a 0.20318s exposure. Non deconvolved images were processed via a macro in ImageJ to sum Z-project, 25 pixel background subtraction (SMA-1:GFP lines), normalize ROI/background area across animals, and mean gray value measured. The difference in intensity was reported here. Microvilli length measurements were performed on the subset of animals with meanders while cilia measurements were obtained via random sampling. Lengths were analyzed with the Simple Neurite Tracer plugin on ImageJ (Longair et al.), with the farthest two points in 3D were assigned manually. All measurements were normalized to body size via pharynx length.

**RNAi**

Overnight *E. coli* HT115 cultures of empty vector, *pros-1* or *hsp-1* RNAi constructs (Ahringer Library)(Kamath, 2003), or *pat-3* RNAi construct (gift from A. Wehman) were used. For *pros-1* and *pat-3* RNAi, gravid adults were bleached (Sulston & Hodgkin, 1988), unsynchronized embryos were placed on seeded RNAi plates, and scored as Day 1 adults. For *hsp-1* RNAi, L4 *rrf-3; nsIs228* animals (Simmer et al., 2002) were picked to RNAi plates and scored 48 hours later. All RNAi experimental data represent ≥2 independent trials.

**Lifespan Assays**

Worms were synchronized via timed egg lay (Wilkinson et al., 2012). Assays were performed in 2+ biological replicates. Since Bag was a prevalent phenotype of *unc-23* mutant animals, data presented here do not censor this, but trends held true also when censored (data not shown).

**AUTHOR CONTRIBUTIONS**

CM and AS designed, performed, and analyzed all experiments, along with JB. AS co-wrote the manuscript with CM, along with JB.

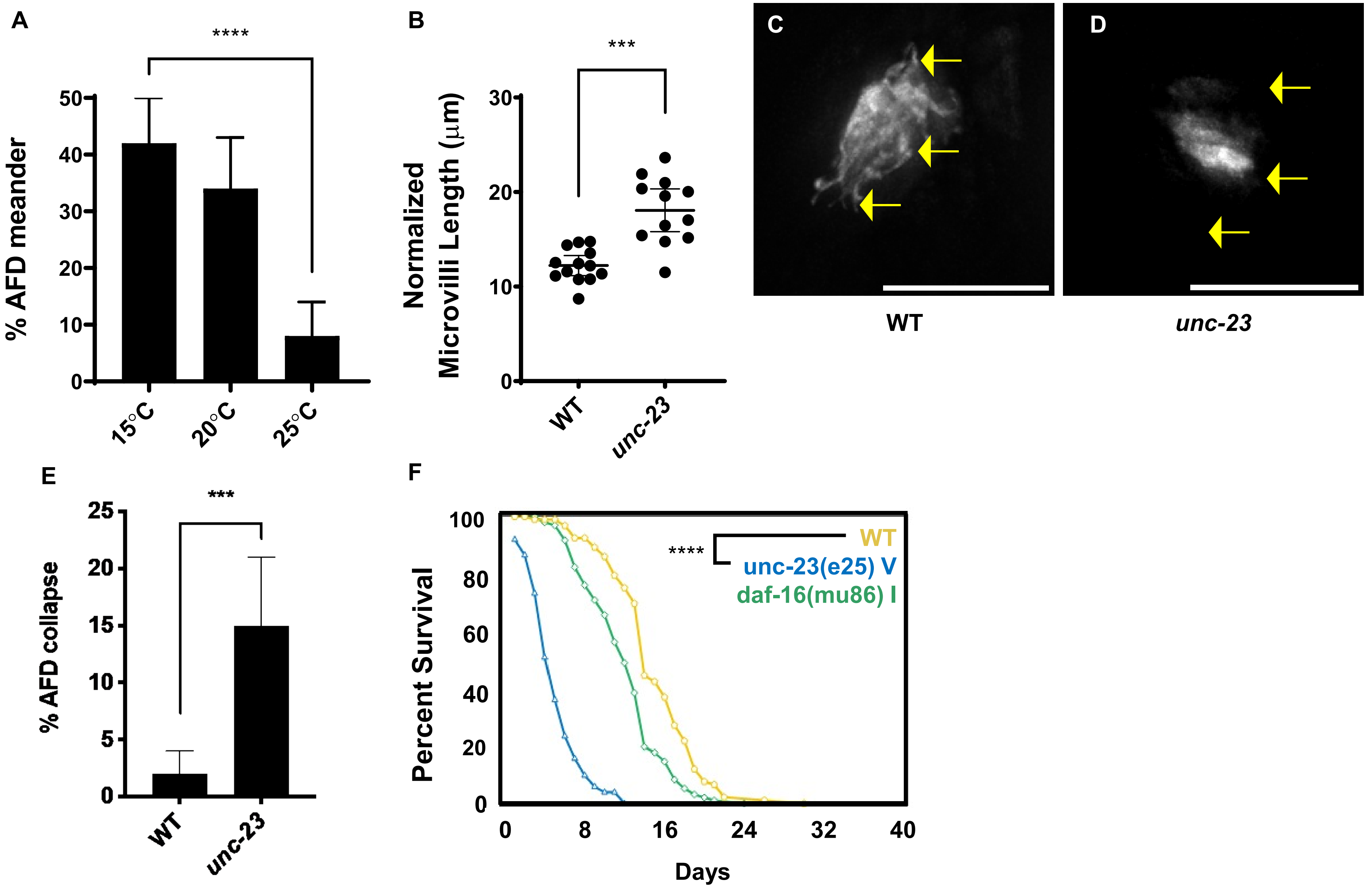
A
****
% AFD meander
50
40
30
20
10
0
15°C
20°C
25°C
B
***
Normalized Microvilli Length (μm)
30
20
10
0
WT
unc-23
C
WT
D
unc-23
E
***
% AFD collapse
25
20
15
10
5
0
WT
unc-23
F
****
WT
unc-23(e25) V
daf-16(mu86) I
Percent Survival
100
80
60
40
20
0
0
8
16
24
32
40
Days

**Figure S1: AFD-NRE meandering phenotype varies with temperature and age**

Analyses on *unc-23(e25)*. AFD-NRE microvilli visualized with *nsIs645*. (**A**) Temperature dependence of % AFD-NRE meander. N≥98 for all conditions. Data plotted as population sum ± 95% CI. Two-sided Fisher's Exact Test **** p<0.0001. (**B**) Meander subset AFD-NRE-m length (µm) for WT (mean= 12.23 ± 1.08 95% CI, N=13) and *unc-23* (mean= 18.07 ± 2.25 95% CI, N=12), normalized to animal size. Data plotted as mean ± 95% CI. Welch's t-test *** p=0.0001. (**C, D**) Fluorescence images of WT (C) and *unc-23(e25)* (D) AFD-NRE collapse in D3 Adults. Scale bar, 8µm. Arrows = AFD-NRE-m, lost in *unc-23* (**E**) % AFD-NRE collapse quantified in WT and *unc-23* mutants. N≥112, plotted as population sum ± 95% CI. Two-sided Fisher's Exact Test *** p=0.0003. (**F**) Representative Kaplan-Meyer lifespan curves of WT, *unc-23(e25)*, and *daf-16(mu86)*. Log Rank (Mantel-Cox) test **** p<0.0001.

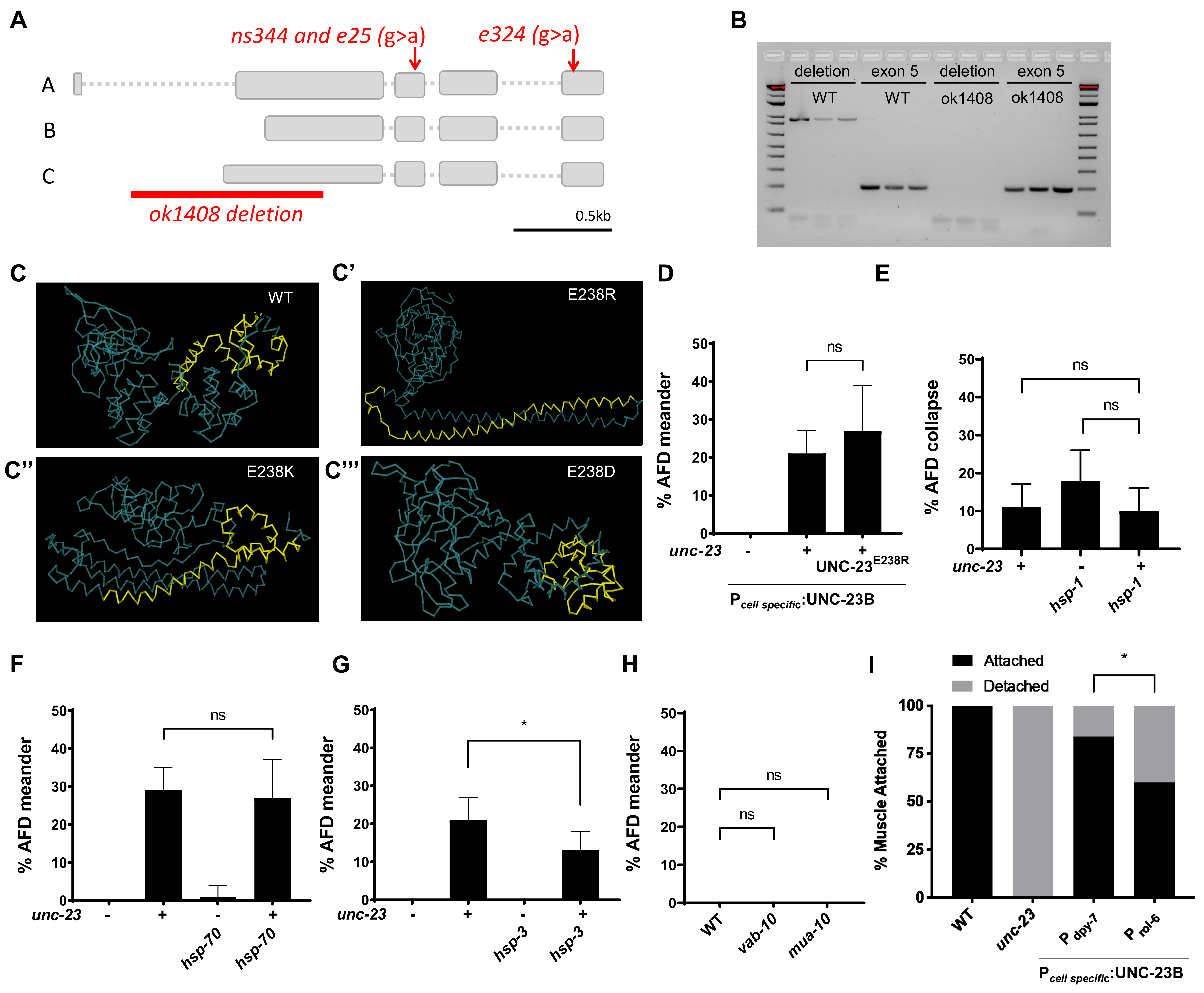
A
ns344 and e25 (g>a)
e324 (g>a)
A
B
C
ok1408 deletion
0.5kb
B
deletion
WT
exon 5
WT
deletion
ok1408
exon 5
ok1408
C
WT
C'
E238R
C''
E238K
C'''
E238D
D
% AFD meander
ns
unc-23
UNC-23E238R
Pcell specific:UNC-23B
E
% AFD collapse
ns
ns
unc-23
hsp-1
hsp-1
F
% AFD meander
ns
unc-23
hsp-70
hsp-70
G
% AFD meander
*
unc-23
hsp-3
hsp-3
H
% AFD meander
ns
ns
WT
vab-10
mua-10
I
Attached
Detached
*
% Muscle Attached
WT
unc-23
P dpy-7
P rol-6
Pcell specific:UNC-23B

**Figure S2: Epithelial UNC-23$^{E238R}$ mutation causes AFD-NRE and muscle defects**

(**A**) Gene structure of *unc-23* with lesions and transcribed isoforms noted. (**B**) Representative RT PCR analysis of WT and *unc-23(ok1408)* shows that the deletion mutant does not transcribe exon 2 but does transcribe exon 5. (**C-C'''**) Phyre2 Intensive protein folding predictions of UNC-23, UNC-23$^{E238R}$, UNC-23$^{E238K}$, and UNC-23$^{E238D}$ as noted. BAG domain = yellow. (**D**) (B) % AFD-NRE microvilli meander in WT (N=201), *unc-23* (N=243), and UNC-23$^{E238R}$ expressing transgenic strain (N=52). Data plotted as population sum ± 95% CI. Two-sided Fisher's Exact Test ns p=0.4631. (**E-G**) % AFD-NRE microvilli meander plotted as population sum ± 95% CI in: (E) *hsp-70(tm2318)* visualized with *nsIs373*. N≥69 except for WT (N=30). Two-sided Fisher's Exact Test ns p=0.7618. (F) *hsp-3(tm832)* visualized with *nsIs228*. N≥195 except for *hsp-3* (N=65). Two-sided Fisher's Exact Test * p=0.0326. (G) WT (N=201), *vab-10(e698)* (N=110), and *mua-10(rh267)* (N=69) visualized with *nsIs228* and plotted as population sum ± 95% CI. Two-sided Fisher's Exact Test ns p>0.9999. (I) % anterior muscle attachment in WT (N= 16), *unc-23* (N= 9), $P_{dpy\text{-}7:}$ UNC-23B; *unc-23* (N=31), and $P_{rol\text{-}6}$:UNC-23B; *unc-23* (N=30) at Day 3 adulthood. Two-sided Fisher's Exact Test * p= 0.0486.

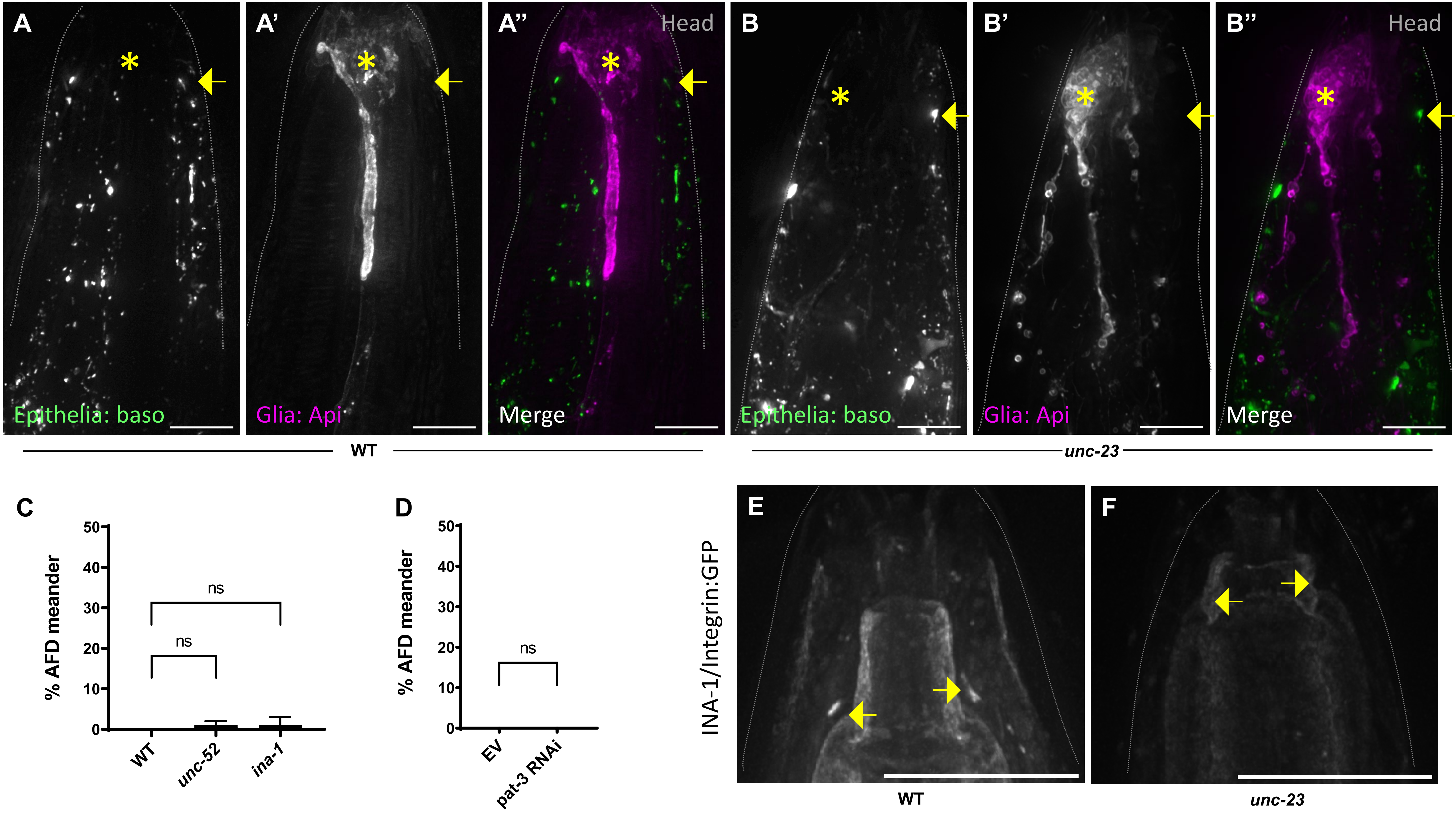
A
A'
A''
Head
B
B'
B''
Head
Epithelia: baso
Glia: Api
Merge
Epithelia: baso
Glia: Api
Merge
WT
unc-23
C
% AFD meander
50
40
30
20
10
0
ns
ns
WT
unc-52
ina-1
D
% AFD meander
50
40
30
20
10
0
ns
EV
pat-3 RNAi
INA-1/Integrin:GFP
E
F
WT
unc-23

**Figure S3: UNC-23 registers Epithelia and AMsh register cell-polarity**

(**A-B''**) Fluorescence images of epithelial basolateral domain marker (green) and AMsh glial apical domain marker (magenta). Merge shows glial apical region (arrow), do not overlap even in overgrowth zones (arrow) *unc-23(e25)* (B-B''). Scale bar, 8µm. (**C**) % AFD-NRE microvilli meander visualized with *nsIs645* and plotted as population sum ± 95% CI in: (A) WT (N=112), *unc-52(e444*) (N=122), and *ina-1(gm144)* (N=87). Two-sided Fisher's Exact Test ns p>0.9999 and p=0.4372. (D) % AFD-NRE microvilli meander with *pat-3* RNAi knockdown (two replicates). Control empty vector (N=68), *pat-3* RNAi (N=32). Two-sided Fisher's Exact Test ns p>0.9999. (**E-F**) Fluorescence images of anterior head region INA-1:GFP (yellow arrows) in anterior head (dotted gray line), lost in *unc-23* mutants (F). Scale bar, 8µm.

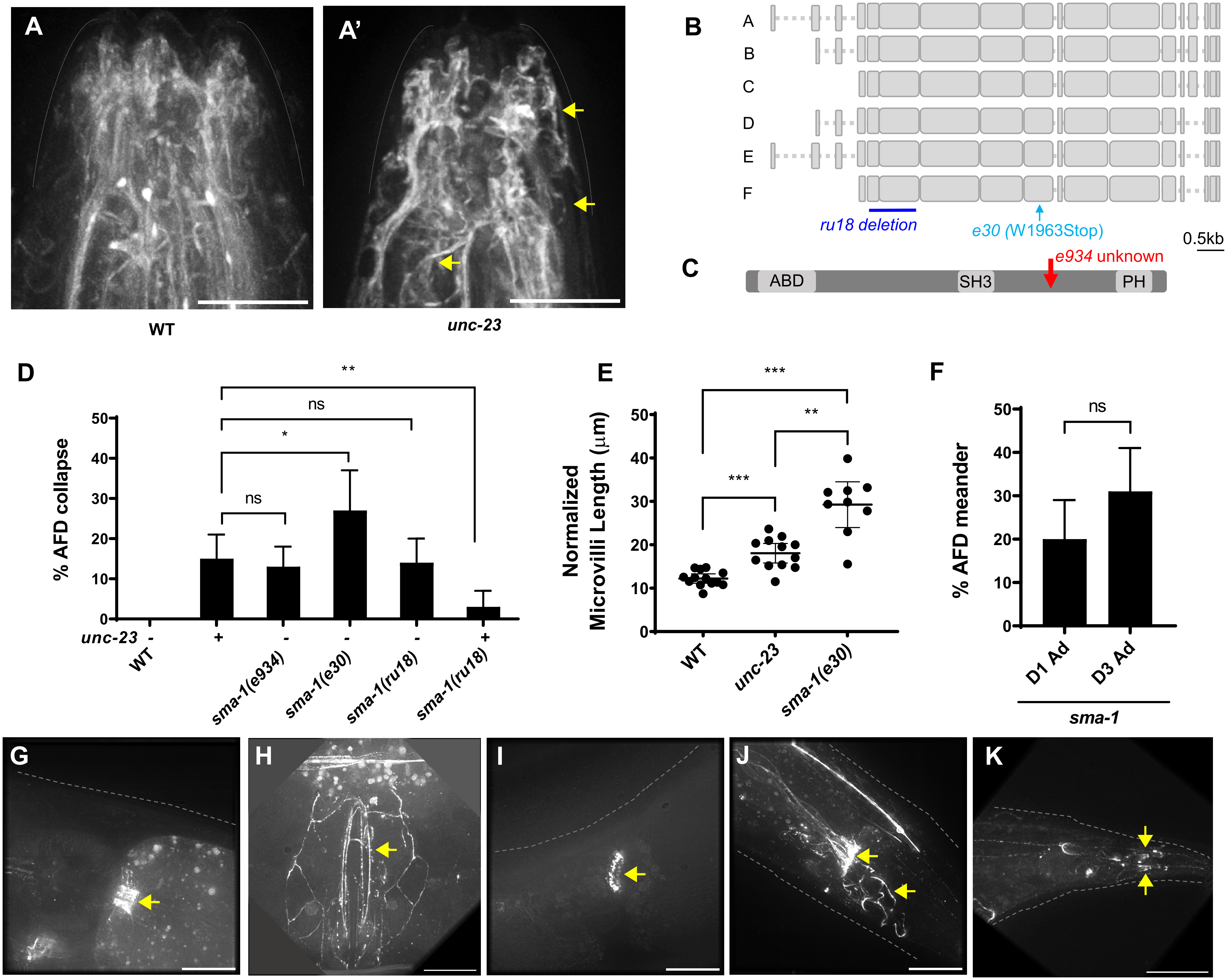
A
A'
WT
unc-23
B
A
B
C
D
E
F
ru18 deletion
e30 (W1963Stop)
0.5kb
C
ABD
SH3
PH
e934 unknown
D
% AFD collapse
unc-23
WT
sma-1(e934)
sma-1(e30)
sma-1(ru18)
sma-1(ru18)
ns
*
**
E
Normalized Microvilli Length (μm)
WT
unc-23
sma-1(e30)
***
**
F
% AFD meander
D1 Ad
D3 Ad
sma-1
ns
G
H
I
J
K

**Figure S4: UNC-23 acts via SMA-1/$\beta_H$ Spectrin and impacts SPC-1/α-Spectrin organization** (**A-A'**) Fluorescence images of SPC-1:GFP in WT (A) and *unc-23(e25)* (A'). Scale bar, 8µm. Dotted line: animal head, anterior nose-tip is top. (**B-C**) Schematic of *sma-1* gene region with lesions noted (B) and SMA-1 protein domains (C) ABD= Actin Binding Domain, SH3= Src Homology Domain, and PH=Pleckstrin Homology domain (**D**) % AFD-NRE collapse phenotype in *sma-1* alleles. N≥88 for all genotypes, data plotted as population sum ± 95% CI. Two-sided Fisher's Exact Test ns p=0.7005 (*e25*-*e934*), * p=0.0352, ns p>0.9999 (*e25*-*ru18*), and ** p=0.0046. (**E**) AFD microvilli length (µm) of WT (mean= 12.23 ± 1.08 95% CI, N=13), *unc-23* (mean= 18.07 ± 2.25 95% CI, N=12), and *sma-1* (mean=29.25 ± 5.27 95% CI, N=9) normalized to animal size and plotted as mean ± 95% CI. Dunnett's T3 Multiple Comparisons Test *** p=0.0003 (WT-*e25*), *** p=0.0001 (WT-*e30*), ** P=0.0028. (**F)** Age dependence of *sma-1(e30)* % AFD-NRE microvilli meander. D1 Ad (N=88) and D3 Ad (N=83) visualized with *nsIs645*, where Ad= adult and D= day. Data plotted as population sum ± 95% CI. Two-sided Fisher's Exact Test ns p=0.1175. (**G-K**) Representative images of SMA-1:GFP expression in an adult animal at the (G) the pharyngeal-intestinal valve, (H) vulva (I) spermatheca-uterine valve, (J) rectum and rectal valve, and (K) distal tail, at presumptive PHsh glial tips. Yellow arrows point to the anatomy noted in each. Scale bar, 15µm.

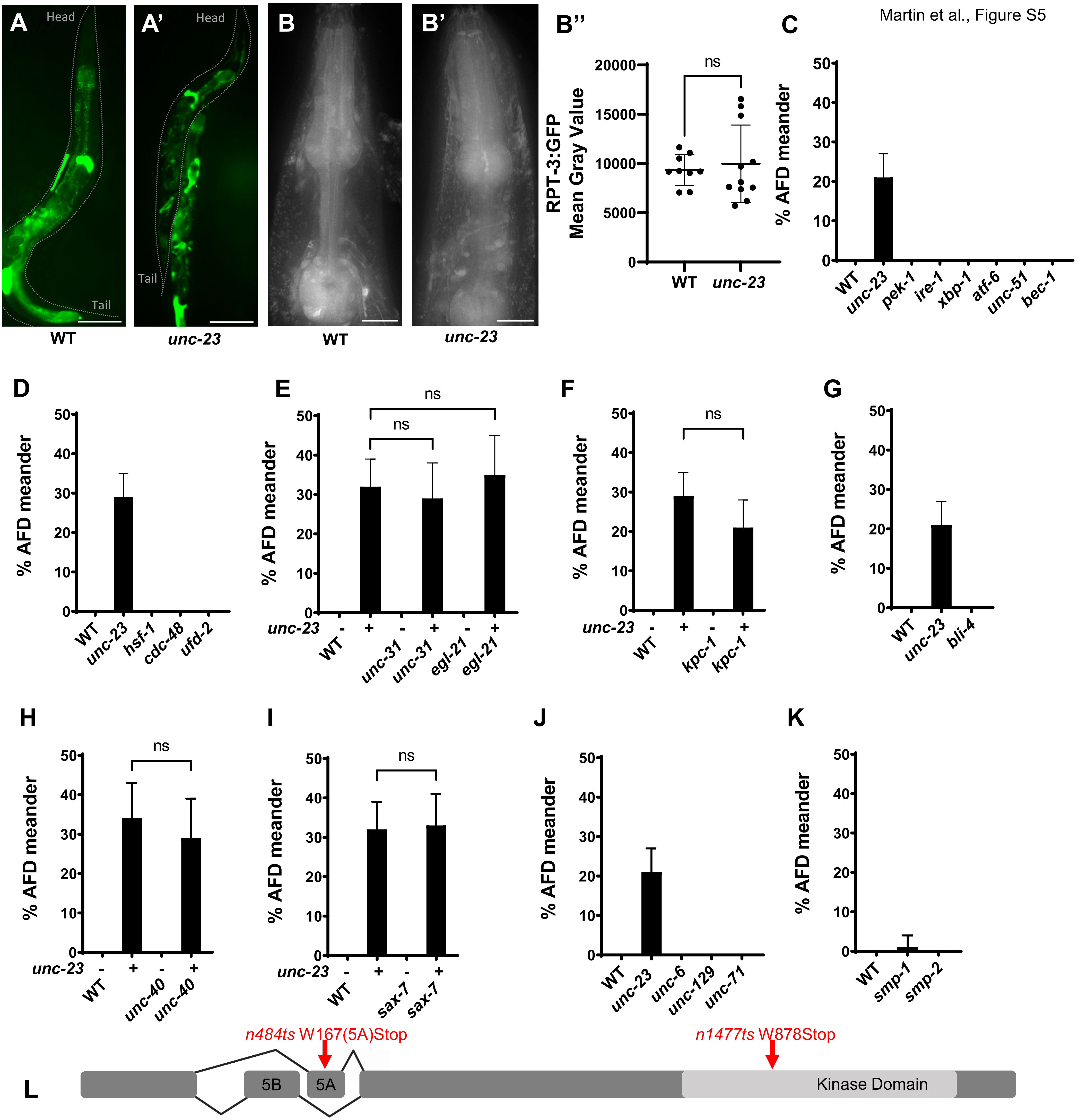
A
Head
Tail
WT
A'
Head
Tail
unc-23
B
WT
B'
unc-23
B''
RPT-3:GFP Mean Gray Value
ns
WT
unc-23
C
% AFD meander
WT
unc-23
pek-1
ire-1
xbp-1
atf-6
unc-51
bec-1
D
WT
unc-23
hsf-1
cdc-48
ufd-2
E
ns
unc-23
WT
unc-31
egl-21
F
kpc-1
G
bli-4
H
unc-40
I
sax-7
J
unc-6
unc-129
unc-71
K
smp-1
smp-2
L
n484ts W167(5A)Stop
n1477ts W878Stop
5B
5A
Kinase Domain

**Figure S5: UNC-23 does not act through general proteotoxicity or stress response pathways**

(**A-A'**) Fluorescence images of $P_{hsp\text{-}4}$:GFP in WT (A) and *unc-23* (A'). Scale bar, 50µm Single animal outlined in each panel. (**B-B''**). Fluorescence images of $P_{rpt\text{-}3}$:GFP in WT (B) and *unc-23* (B'). Scale bar, 15µm. Animal head, dotted outline. RPT-3:GFP Mean Gray Values quantified in (B''). WT (N=9), *unc-23* (N=11). Two-sided Welch's t-test ns, p=0.6329 (**C-L**) Data plotted as population sum ± 95% CI. % AFD meander in: **(C)** WT (N=201), *unc-23(e25)* (N=243), *pek-1(ok275)* (N=78), *ire-1(zc14)* (N=112), *xbp-1(zc12)* (N=136), *atf-6(ok551)* (N=64), *unc-51(e369)* (N=34), and *bec-1(ok700)/nT1[qIs51]* (N=68) visualized with *nsIs228*. (D) WT (N=30), *unc-23(e25)* (N=177), *hsf-1(sy441)* (N=130), *cdc-48(tm544)* (N=54), and *ufd-2(tm1380)* (N=78) visualized with *nsIs373*. (E) WT (N=201), *unc-23(e25)* (N=170), *unc-31(e928)* (N=125), *unc-31(e928); unc-23(e25)* (N=94), *egl-21(n476)* (N=99), and *egl-21(n476); unc-23(e25)* (N=85) visualized with *nsIs228*. Two-sided Fisher's Exact Test ns p=0.5805 (*e25- e25; e928*) and p=0.6736 (*e25- e25; n476*). (F) WT (N=30), *unc-23(e25)* (N=177), *kpc-1(gk8)* (N=75), and *kpc-1(gk8); unc-23(e25)* (N=102) visualized with *nsIs373*. Two-sided Fisher's Exact Test ns p=0.1558. (G) WT (N=201), *unc-23(e25)* (N=243), *bli-4(e937)* (N=116) visualized with *nsIs228*. (H) WT (N=112), *unc-23(e25)* (N=122), *unc-40(e271)* (N=121), and *unc-40(e271); unc-23(e25)* (N=92) visualized with *nsIs645*. Two-sided Fisher's Exact Test ns p=0.4628. (I) WT (N=201), *unc-23(e25)* (N=170), *sax-7(ky146)* (N=130), and *sax-7(ky146); unc-23(e25)* (N=128) visualized with *nsIs228*. Two-sided Fisher's Exact Test ns p>0.9999. (J) WT (N=201), *unc-23(e25)* (N=243), *unc-6(e78)* (N=54), *unc-129(ev557)* (N=71), and *unc-71(e541)* (N=72) visualized with *nsIs228*. (K) WT (N=72), *smp-1(ev715)* (N=72), and *smp-2(ev709)* (N=61). (**L**) *egl-15* gene region showing lesions and alternate splicing of exon 5 to generate A or B isoforms.

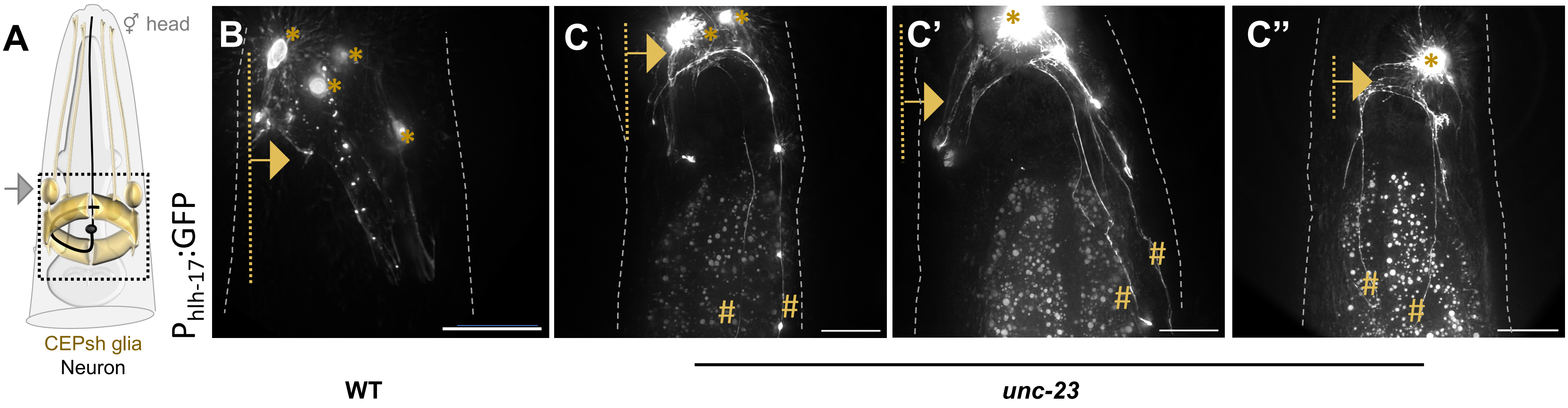
A
⚥ head
CEPsh glia
Neuron
B
C
C'
C''
$P_{hlh-17}$::GFP
WT
unc-23

**Figure S6: CEPsh glia posterior sheath processes are defective in *unc-23(e25)* mutants**

(**A**) Schematic of CEPsh glia in animal head, dashed box denotes region imaged in B-C''(**B-C''**) Fluorescence images of CEPsh glia posterior processes enwrapping the brain neuropil in WT (B) and *unc-23* (C-C''). Scale bar, 15µm. Gray outlines animal in each panel. Dotted yellow line and arrow notes normal anatomical domain of CEPsh posterior sheaths, hash yellow points to aberrant extension of these.
